# Contribution of the Golgi apparatus in morphogenesis of a virus induced cytopathic vacuolar system

**DOI:** 10.1101/2022.03.14.484265

**Authors:** Ranjan Sengupta, Elaine M. Mihelc, Stephanie Angel, Jason K. Lanman, Richard J. Kuhn, Robert V. Stahelin

**Affiliations:** Department of Medicinal Chemistry and Molecular Pharmacology, Immunology and Infectious Disease, Purdue University, West Lafayette, IN 47907, USA; Department of Biological Sciences, Immunology and Infectious Disease, Purdue University, West Lafayette, IN 47907, USA; The Purdue Institute of Inflammation, Immunology and Infectious Disease, Purdue University, West Lafayette, IN 47907, USA

**Keywords:** alphavirus, cytopathic vacuoles, electron tomography, Golgi remodeling, membrane bending, transmission electron microscopy

## Abstract

The Golgi apparatus (GA) in mammalian cells is pericentrosomally anchored and exhibits a stacked architecture. During infections by members of the alphavirus genus, the host cell GA is thought to give rise to distinct mobile pleomorphic vacuoles known as CPV-II (cytopathic vesicle-II) via unknown morphological steps. To dissect this, we adopted a phased electron tomography approach to image multiple overlapping volumes of a cell infected with Venezuelan equine encephalitis virus (VEEV) and complemented it with localization of peroxidase tagged Golgi marker. Analysis of the tomograms revealed a pattern of progressive cisternal bending into double-lamellar vesicles as a central process underpinning the biogenesis and the morphological complexity of this vacuolar system. Here we propose a model for the conversion of GA to CPV-II that reveals a unique pathway of intracellular virus envelopment. Our result has implications for alphavirus virus induced displacement of Golgi cisternae to the plasma membrane to aid viral egress operating late in the infection cycle.

## Introduction

In eukaryotic cells, membrane-bound organelles transition over their lifetime through multiple morpho-functional states (Heald and Cohen-Fix, 2014; Westrate et al, 2015). In addition to endogenous events, morphological remodeling of organelles is also observed during virus infection of the host cell. Viruses have evolved to exploit host cell structural and functional plasticity of organelles and thus are important tools to dissect and understand cellular trafficking pathways (Den Boon et al, 2010; Chen et al, 2015; Cortese et al, 2017; Hollinshead et al, 2012; Hsu et al, 2010; Romero-Brey et al, 2012; Welsch et al, 2009). Indeed, a major restructuring of the host-cell secretory system has been observed associated with virus maturation and egress leading to either delayed or little consequences on cellular secretion (Mousnier et al., 2014); an indication that these pathogens operate within the critical structural and functional parameters of host organelle function. These recurrent observations necessitate the investigation of pathogen induced structural states of host organelles in order to obtain clues on morpho-functional states observed during viral infection.

The genus alphavirus, from the *Togavirdae* family are enveloped, icosahedral (∼700 Å diameter), positive-strand RNA viruses that include significant human pathogens such as Chikungunya virus (CHIKV) and Venezuelan equine encephalitis virus (VEEV). Their genomic RNA is packaged by capsid protein to form a spherical nucleocapsid core (NC) (∼40nm in diameter) (Jose et al., 2009; Zhang et al., 2011). Typically, the NC obtains its envelope via budding at the plasma membrane facilitated by the interaction of the membrane embedded envelope glycoproteins E1/E2 and capsid (Brown et al, 2018). The glycoproteins assemble in the ER, undergo maturation in the Golgi apparatus (GA) and are then transported to plasma membrane (PM) where they interact with capsid to initiate budding (Sanchez-San Martin et al, 2009). However, during mid to late stages of the alphavirus infection cycle, a structurally distinct pleomorphic vacuolar system (cytopathic vesicles II or CPV-II) predominates in the cytoplasm of the host cell. CPV-II are known to originate from the GA and exhibit a characteristic accumulation of nucleocapsid core (NC) on its membrane (Grimley et al, 1968; Mussgay and Weibel, 1962). These early studies revealed the morphological complexity of CPV-II where both uni- and bi-lamellar forms of CPV-II were observed in the milieu. Griffiths et al. demonstrated that monensin blocked E1/E2 glycoprotein transport out of the GA and resulted in the swelling and vacuolization of Golgi cisterna that bound nucleocapsid cores (much like the CPV-II) as a result of accumulation of viral glycoprotein in the GA (Griffiths et al, 1983). Although this mechanism could explain biogenesis of the unilamellar CPV-II, it does not account for other forms of CPV-II, especially the bi-lamellar form reported in the early studies. Additionally, from a cell biology perspective, neither structural nor the functional consequence of this vacuolar biogenesis on the cell secretory system is exactly known.

The structure of the GA is known to be intrinsically connected to its various functions. During interphase, the mammalian GA is pericentrosomally anchored and exhibits a stacked structure comprised of a functionally defined set of cisternae (namely, cis-, medial- and trans) (Klumperman, 2011; Yadav and Linstedt, 2011). However, at instances such as cell division or during certain bacterial and viral infections, the GA loses its typical stacked architecture and its pericentrosomal localization (Nakagomi et al, 2008; Petrosyan et al, 2014; Thayer et al, 2013; Wei and Seemann, 2017; Campadelli et al, 1993; Gahmberg et al, 1986; Hansen et al, 2017; Heuer et al, 2009; Kim and Satchell, 2016; Lavi et al, 1996; Quiner and Jackson, 2010). Though understandably critical to its function, a clear morphological understanding of the Golgi membrane remodeling pathways and their exact functional relevance is still lacking. Thus, such instances of virus induced morphological remodeling of the GA could provide excellent opportunities for studying the morpho-functional plasticity of this secretory organelle.

Here we employed innovative transmission electron microscopy (TEM) approaches to elucidate host-cell GA conversion into various forms of CPV-II during infection of TC-83, the vaccine strain of VEEV. Using traditional thin-section TEM, we established a timeline of structural changes of the GA during infection. We then applied an improvised hybrid sample fixation method in conjunction with peroxide tagging of a GA marker to show that previously identified forms of CPV-II carry this marker. To gain a three-dimensional (3D) understanding of the process we established a phased data collection approach and took advantage of the stability of resin sections under the electron beam to collect multiple tilt-series from the same area of the cell over time. The vast array of 3D data collected enabled statistical analyses that yielded a 3D classification of the CPV-II system, identification of large and complex intermediates that links the various forms of CPV-II, the relative spatial frequency of the various forms, and a model of their morphogenesis. Thus, our data identifies the GA-associated pathway of CPV-II biogenesis and proposes a model for the restructuring of Golgi cisternae into large single and double lamellar CPV-II forms identified in VEEV infection. Thus, this work outlines a model for the morphogenesis of virus induced structures from the GA that has implications for an accessory egress mechanism in late stages of the virus life cycle.

## Results

To carry out this work we chose to use the live-attenuated vaccine strain of VEEV, TC-83. TC-83 can be handled in BSL-2, making it possible to use live infected cells for high-pressure freezing (stationed within BSL-2 lab) to provide high quality membrane preservation required for such studies. We ruled out the possibility that the critical mutation present on E2 (T120R) could influence the binding of NC to E2, the focus of this study, by modelling the mutation to the local structure of E2 (Fig S1).

### VEEV infected cells show progressive remodeling of the GA with and emergence of pleomorphic cytopathic vesicles

A considerable GA remodeling precedes the onset of CPV-II. Golgi structures in BHK cells were screened at different time points starting at 3 hours postinfection (PI) up to 12 hours (Fig S2, A-E). These features on the GA were absent in uninfected BHK cells (Fig S3, red arrows). The GA architecture at 12 hours PI exhibited extensive herniations and vacuolization in addition to unstacking and bending (Fig S2B, i-iv). 250 nm serial sections were screened for viewing large complex structures (Fig S2C, i, sections 1 through 20). The pleomorphic nature of the CPV-II was more evident in these thicker sections as oblong and dumb-bell shaped structures studded with NC were readily observed. At 12 hours PI, multiple CPV-II clusters were observed near the plasma membrane that were composed of various morphological forms of CPV-II (Fig S2E, i to iii). At this stage, CPV-II exhibited a more scattered distribution in the cytoplasm.

To detect the distribution of both early and late vacuoles (CPV-II) arising from the GA, we carried out HRP-tagging based localization of the Golgi marker α-mannosidase-II. Here we employed a recently published hybrid method for superior sample preservation to localize HRP-tagged α-mannosidase-II (Golgi FLIPPER, Kuipers et al, 2015), a cis-medial Golgi marker (Sengupta et al, 2019). BHK cells first transfected with ManII-HRP encoding plasmid were then infected with TC-83 virus (MOI of 20) and then fixed at 6 and 12 hours PI.

In control cells, a tight perinuclear staining was observed. At a higher magnification, the perinuclear stain resolved into multiple stacks of GA (Fig 1A ii and iii, blue arrowheads and area within blue box) (Fig 1A i, region demarcated with red box). In contrast, infected samples from 6 hours PI showed stained vesicular structures scattered predominantly in the perinuclear region (Fig 1B i, region demarcated with yellow box). At higher magnifications they resolved into multiple stained Golgi stacks (Fig 1B iii-v, blue arrowheads), and large vesicles (∼200-500 nm) associated with the stacks (Fig 1B iii, yellow arrows). Further, large cisternal herniations gave rise to distended structures that were still connected to the Golgi stack (white and magenta arrows, Fig 1B iii to v). Distended structures originated from both the outer periphery of the stacks (Fig 1B iii and iv, white arrows,) as well as from the middle of the stack (magenta arrows, Fig 1B iv and v). A limited number of isolated vesicles carrying the ManII-HRP marker were detected near the plasma membrane (PM) (Fig 1B vi, green arrows). The samples imaged 12 hours PI revealed CPV-II carrying the ManII-HRP stain at a high concentration not only at the perinuclear regions but also scattered throughout the cytoplasm and near the plasma membrane (area demarcated by orange box in Fig 1C i and S3A i through vi). Images of the representative section show stained vesicular structures at the perinuclear region (orange box in Fig 1C i). Images of this region at a high magnification exhibit clusters of CPV-II (Fig 1C ii, denoted by red arrows) and a mix of various morphological forms (Fig 1C iii, indicated with roman numerals I, II or IV and their tomographic classification in Fig 6). A dense complement of NC was observed associated with the cytopathic vesicles in these clusters.

**Figure 1.**
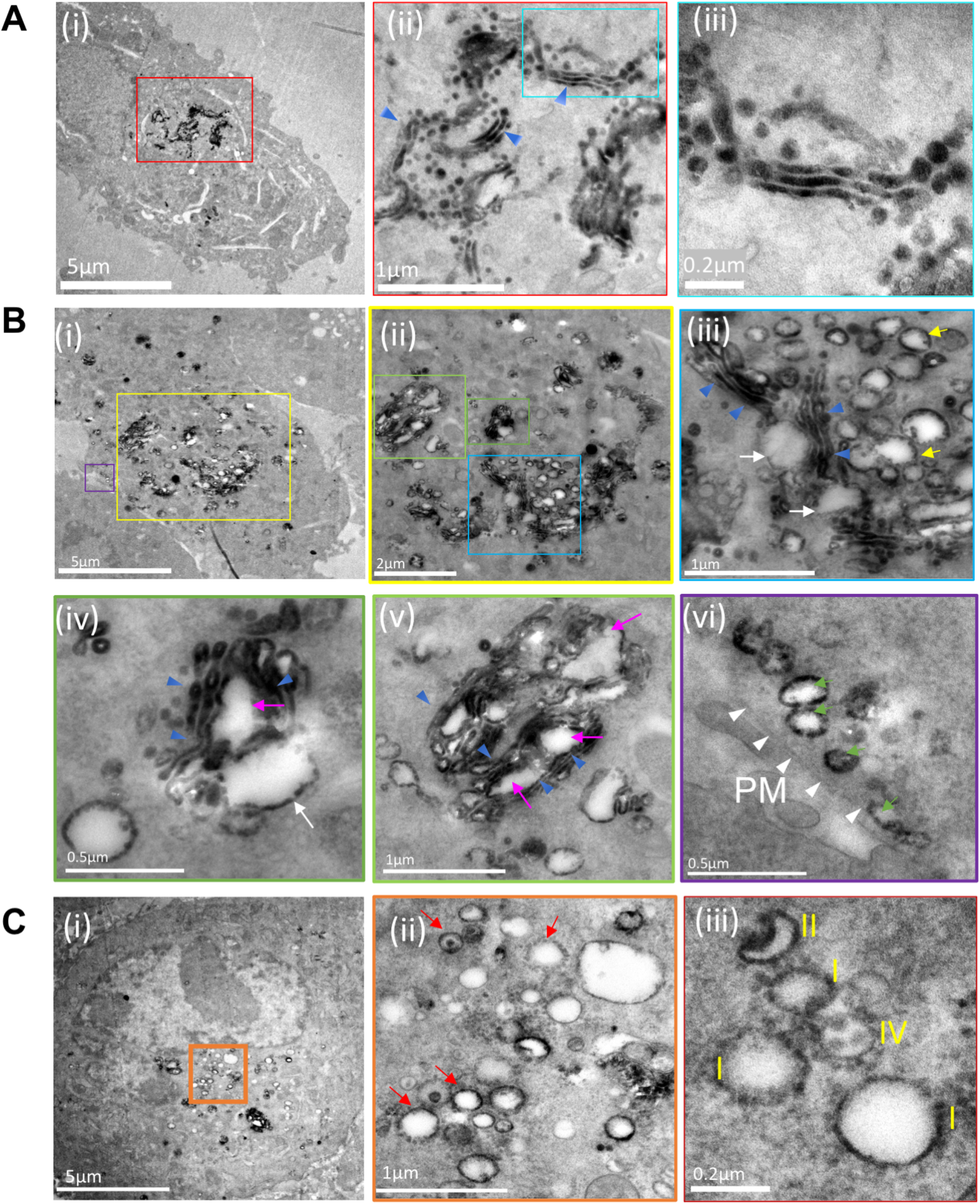
Pleomorphic CPV-II in the perinuclear region and near the plasma membrane are of Golgi origin. A hybrid chemical-cryo fixation method was optimized with HRP-DAB assay for the detection of HRP-tagged α-mannosidase-II, a (cis-medial) Golgi membrane marker in mock infected and in VEEV infected BHK cells at 6 and 12 hours postinfection (PI) (**(A)** (i)-(iii), **(B)** (i)-(vi) and **(C)** (i)-(iii), respectively). TEM images of 90 nm resin sections of mock controls show a tight perinuclear staining (**(A)**(i), area within red box) that at higher magnification resolves into multiple canonical Golgi stacks (**(A)**(ii)). Magnified view of the specifically labelled single stack exhibit well preserved cis-medial stacks (**(A)**(iii)) area magnified within the blue box in (ii). Cells from 6 hours PI show the manII-HRP Golgi marker scattered mostly at the perinuclear region with limited punctae outside this region **(B)**(i)] Successive magnified views (**(B)**(ii) through (v)) of the perinuclear region (**(B)**(ii)) demarcated with a yellow box in (i) and color coded henceforth show herniated cisternae within the existing Golgi stacks (**(B)**(ii)-(iv), blue arrow heads), vacuolization (white arrows, (iii)-(v)) and the area around littered with vesicles of similar sizes. These herniations were not restricted either to the *trans-* or the *cis-* face but were seen to occur from cisternae in the middle of the stacks (white arrows, **(B)**(iii)-(v)). Progressive conversion of the cisternae into large vesicles and the replacement of a stacked structure with a cluster of vesicles was apparent **(B)**(iv)). These Golgi derived vesicles (**(B)**(vi), yellow arrows) were also found in limited numbers at the plasma membrane ((vi), demarcated with white arrows). **(C)** Images of cells sections here from 12 hours PI exhibit different classes of CPV-II (**(C)**(i), demarcated with a red box) and show the absence of stacked Golgi structures but a predominance of CPV-II with detectable nucleocapsid cores (NC) associated with the vesicles (ii, red arrows). Some NCs were also detected in free space within these clusters (iv, white arrowheads), but could not be concluded if they were freely occurring in the cytoplasm or were attached to CPV-II excluded from the thin section during sectioning. The most prominent classes of vesicular structure observed were namely, a crescent shaped structure with higher density towards the inner curved side indicating the presence of NC, a typical round vesicular structure with NC on the cytoplasmic surface, vesicle with enveloped virus within and NC on the outer surface (**(C)**(ii) and (iii)). See (Fig S3) for distribution of HRP-ManII carrying vesicles at the same time point and a comparison with HeLa cells stably expressing HRP-ManII.

Additionally, in order to determine if this Golgi vacuolization is cell type specific, HeLa cells stably expressing ManII-HRP were infected with TC-83. Results from the screen in HeLa cells at 6 hours PI (Fig S4B) show similar features observed in BHK cells. As with BHK cells, while mock infected cell lines show tight perinuclear localized Golgi stacks (Fig S3B), infected cells exhibit Golgi stacks that were restricted mainly at the perinuclear region with cisternal herniations and vacuolization (Fig S4B iv-vi). This validation from the ManII-HRP-HeLa cell line also rules out the possibility of artefactual localization of the marker and perturbed morphology of the GA as a result of overexpression.

### Serial-section tomographic reconstruction of a representative infected cell

In order to visualize the structural changes in a structurally complex large organelle such as the Golgi, a combination of serial sectioning and image montaging was employed to obtain large volume tomograms. First, 250 nm thick serial sections of the samples were collected and screened for cells containing multiple Golgi stacks exhibiting a structural flux and a presence of abundant CPV-II. A suitable area in a typical representative cell was then identified, the volume of interest within which spanned several consecutive sections on the EM grid (see materials and methods for details). This first volume (tomogram 1, area outlined in cyan in Fig 2B (see also Video S1, segmented tomogram) consisted of 3 well defined Golgi stacks formed the basis of our GA study. Segmentation of this reconstructed volume consisting of the three Golgi stacks revealed a pathway of Golgi conversion into a vesicular system (Fig 2C, Golgi stacks in green and CPV-II in gold). Reconstruction and segmentation of tomogram1 provided clues that led us to the collection of tilt-series for tomogram 2 (Fig 6E(b)) and tomogram 3 (Fig 8) from the same cell.

**Figure 2.**
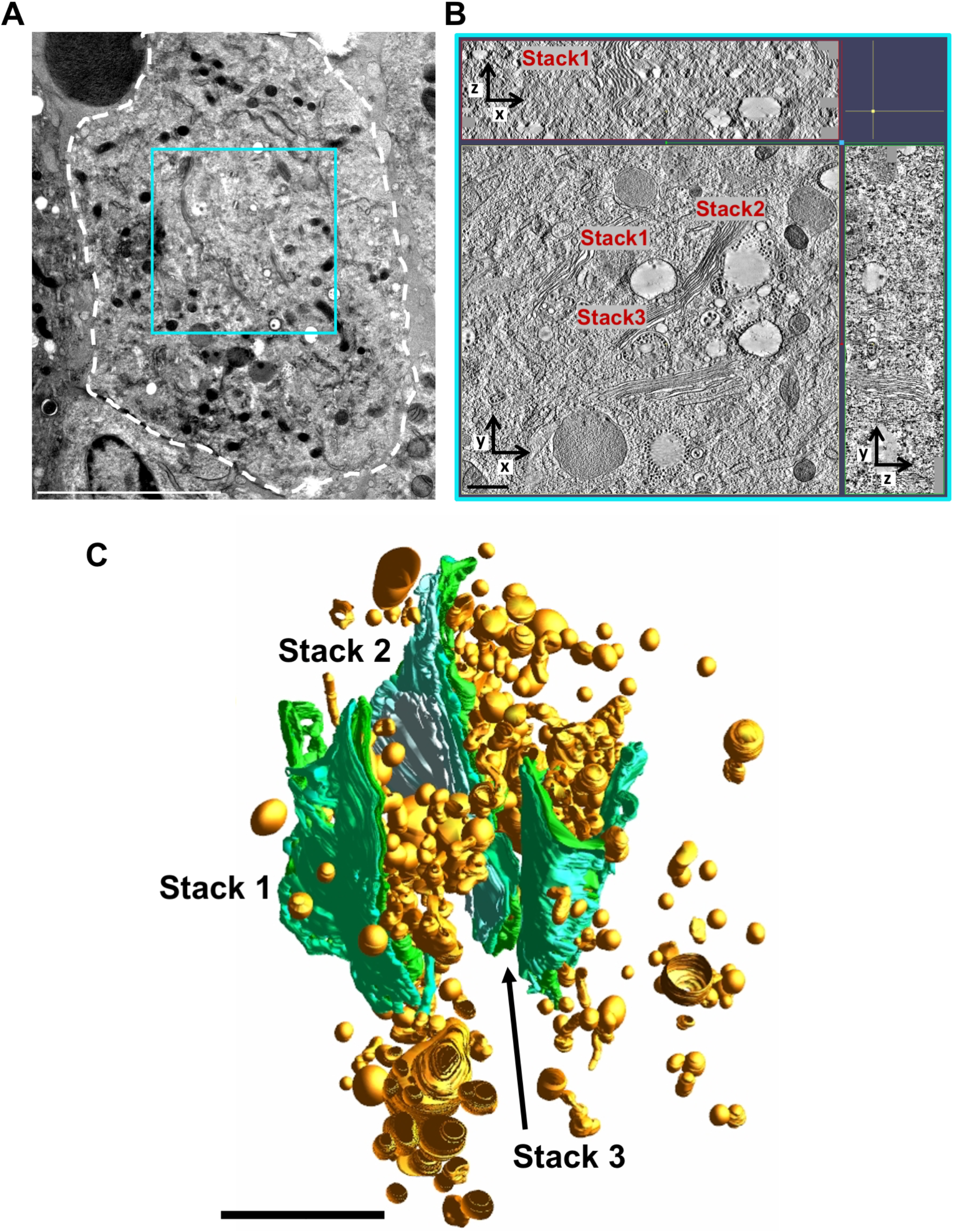
Large volume tomogram of a VEEV-infected cell. **(A)** TEM image collected at 300 kV of a 250-nm thick section of a TC-83-infected BHK cell (outlined in white). The approximate area of this cell from where 8-10 serial sections were used to collect and reconstruct tomogram 1 is demarcated by the cyan box. **(B)** A 3-nm thick virtual section from tomogram 1 **(See also video S1)** with the z-depth along the x- and y-axis shown along the top and right side, respectively. The depicted slice exhibited three Golgi stacks that are denoted by stack 1, 2 and 3 respectively **(C)** 3D visualization of the tomogram volume employing manual segmentation of Golgi (green) and nascent CPV-II Golgi (gold). Scale bars: **(A)** 5 μm, **(B)** 500 nm, and **(C)** 2 μm.

**Figure 3.**
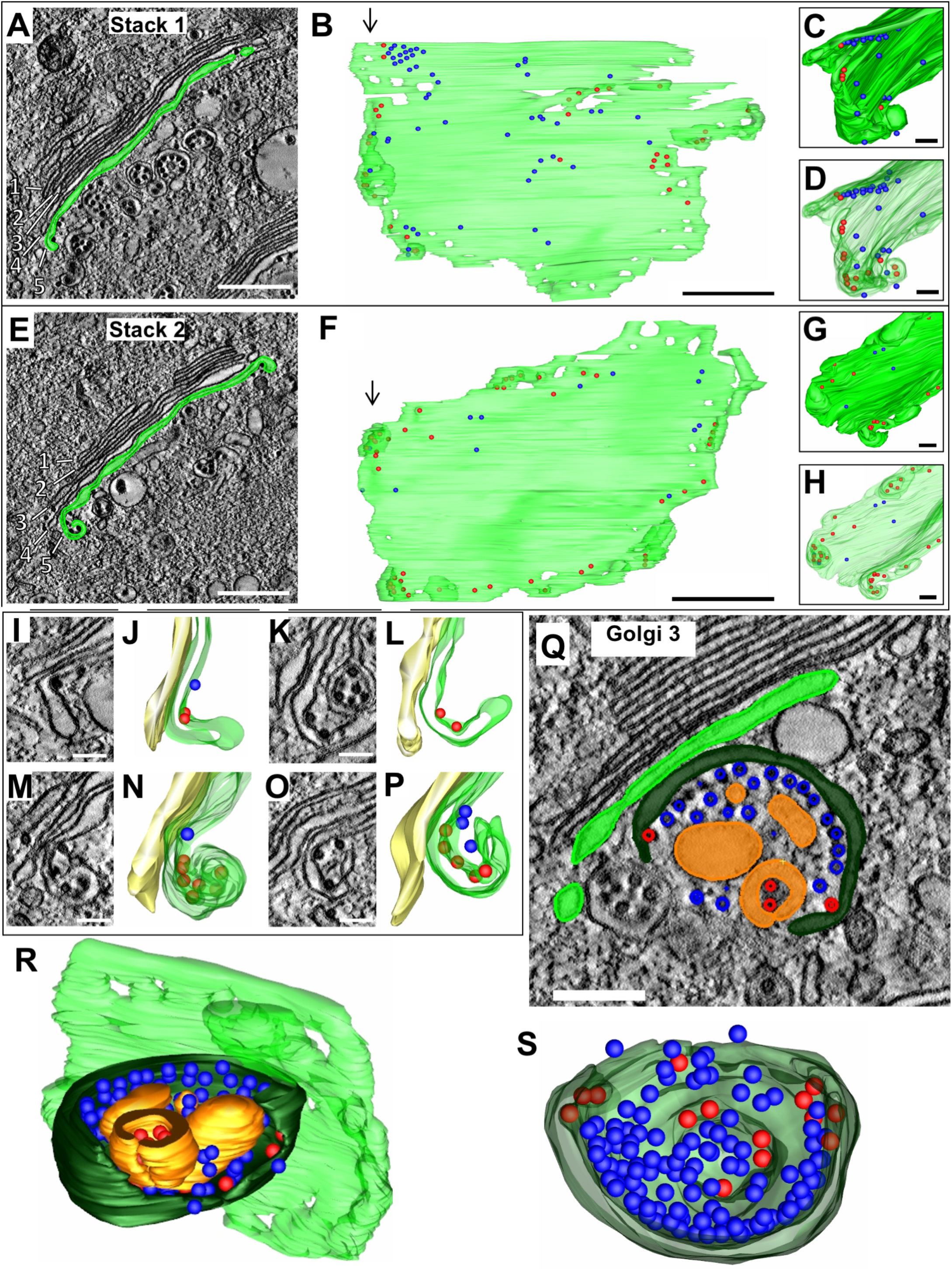
Golgi cisternal rims exhibit bending in regions in contact with NC. **(A)** 3-nm thick virtual section from tomogram 1, showing a Golgi stack (“Stack 1”) with the final intact cisterna colored green. Five cisternae are visible. **(B)** Three-dimensional rendering of the cisterna through the entire 2.5 μm depth of the reconstructed volume. Colored spheres represent NCs in contact with the Golgi membrane, colored blue or red based on the radius of curvature on the adjacent membrane as shown in Fig S6. **(C-D)** The curled edge of the cisterna viewed from the top down the cisternae in the direction of the arrows in panel B. **(E-H)** Similar views as A-D of a second Golgi cisterna through the depth of tomogram 1. **(I-P)** Edges of the cisternae (green) exhibit various degrees of curling away from the next cisterna in the stack (yellow). **(Q-S)** A third Golgi stack with a large portion of a cisterna (dark green) bending away from the adjacent intact cisterna (light green). The tomographic slice **(Q)** and 3D renderings **(R-S)** of this cupped cisterna which is bound with NC (red and blue) and encloses a double-membrane CPV-II (gold). See video **S2 and S3**. Scale bars: 500 nm **(A, E)**, 100 nm **(I, M, K, O)**, and 200 nm **(Q)**.

**Figure 4.**
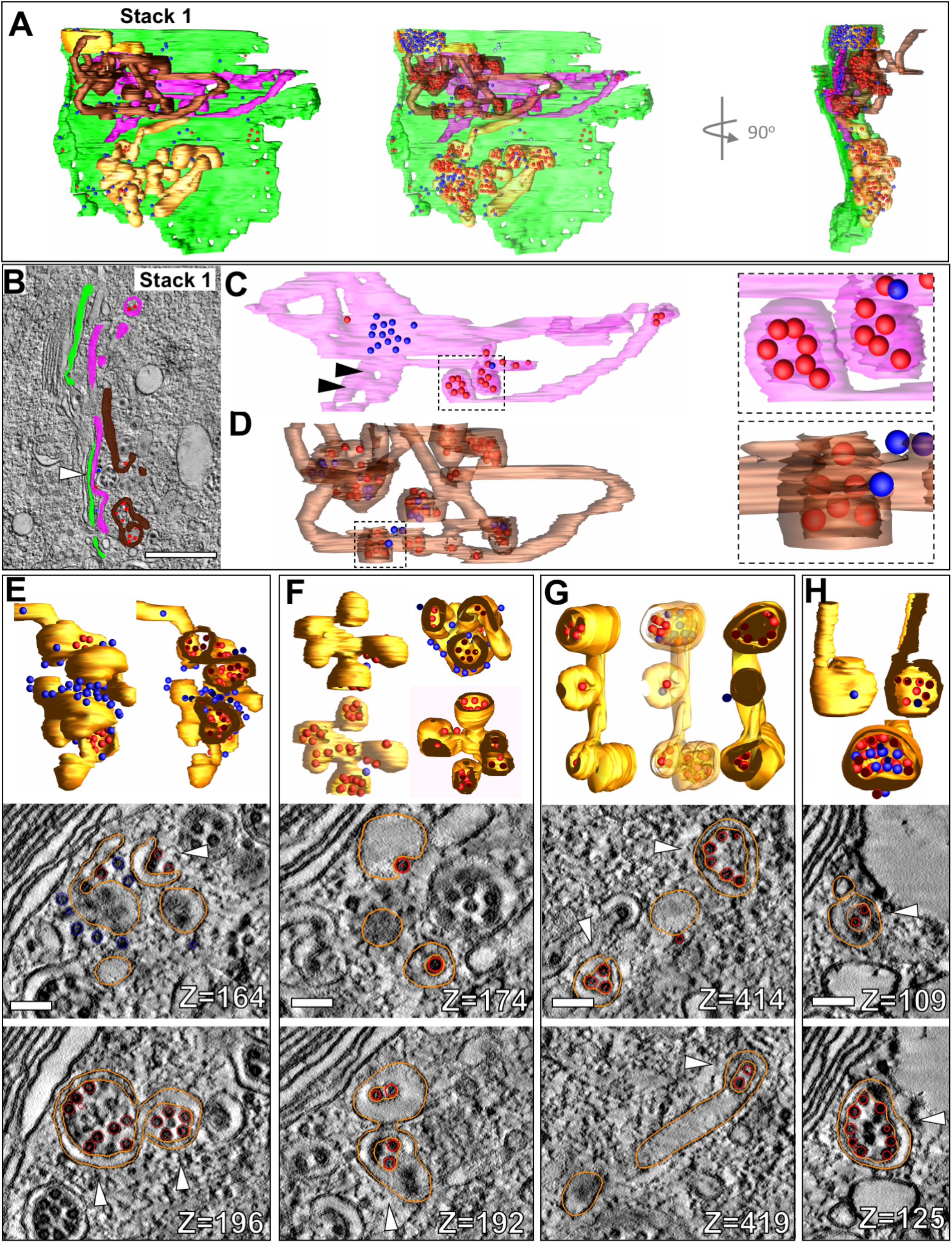
Pleomorphic membrane structures with bound NC are fragmented Golgi cisternae. (A) 3D rendering of Golgi 1 *trans*-most cisterna along with selected pleomorphic CPV-II forms (gold) closest to cisternae (pink and brown) and associated NC (blue and red spheres). **(B)** 3-nm virtual section through stack 1, highlighting the intact cisterna (light green) and two large, fragmented cisternae (magenta and brown) and their associated NC (blue and red spheres). **(C-D)** 3D renderings of two pleomorphic cisternal forms which are highlighted in magenta **(C)** and brown **(D)** in B. Insets of C and D show double-lamellar nature of vesicular structures, with NC inside (red) and outside (blue). CPV-II structures of varying pleomorphic shapes include forms with several double-lamellar vesicular formations **(E-F)**, as well as other vesicular and tubular elements **(G-H)**. Cutaway views in E-H reveal the double-lamellar nature of the vesicular forms. 3-nm thick virtual sections from two different depths of each form shown in E though H are presented with the segmentation of the structures of interest shown in gold and NC in red. The double-lamellar nature of the forms (similar to class 2 and 3) is indicated by white arrowheads. See video S4. Scale bars: 500 nm **(B)** and 100 nm **(E-H)**.

**Figure 5.**
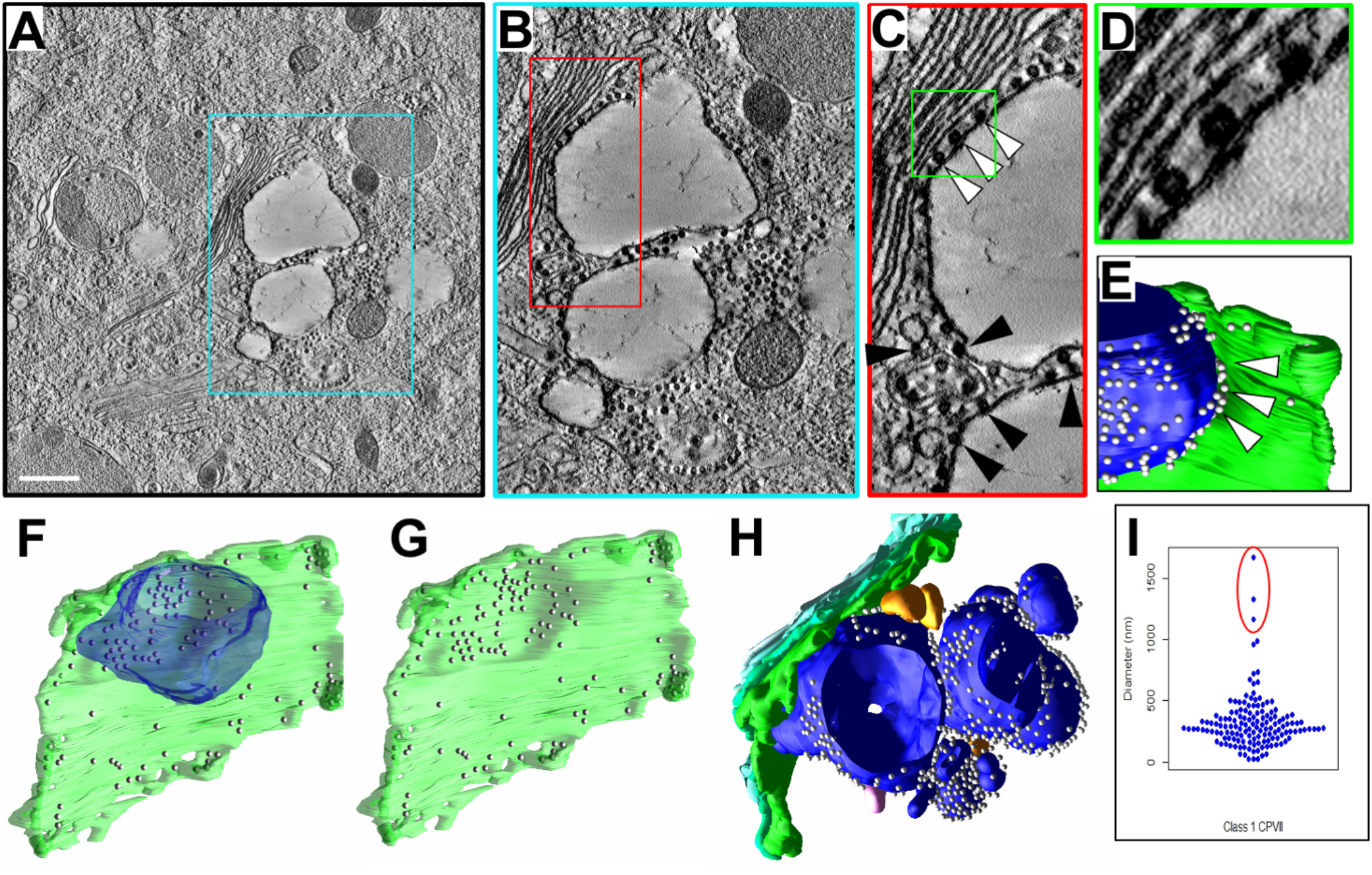
Three-dimensional analysis of Golgi associated vacuoles. **(A-D)** A 3-nm thick virtual section from tomogram 1 showing a large CPV-II abutting the intact *trans*-most cisterna of a Golgi stack, with increasing magnification of the region of interaction presented in the color-coded outlined areas in A-D. NC can be observed making contact with both the intact cisternal membrane and the vesicular membrane (**C**, white arrowheads and **D**), as well as between multiple CPV-II in the cluster (**C**, black arrowheads). **(E)** A 3D view of the segmented CPV-II region shown in C-D (blue), with NC (white spheres) filling much of the gap between the vesicle and the Golgi (green), indicated by arrowheads. **(F-G)** A 3D view depicting the entire *trans-*most cisternae shown in A-E with **(F)** or without **(G)** the large CPV-II. A concentration of NC is seen on the Golgi at the interface with the CPV-II, as well as an indentation in the Golgi where the vesicle was located. **(H)** 3D segmented view of the cluster of CPV-II surrounding the large vesicle depicted in F, consisting of variably sized class 1 CPV-II (blue) and their associated NC, and a few class 3 CPV-II (gold), all in close physical proximity with the Golgi stack (green shades). The three largest class 1 CPV-II in the volume are found in this particular cluster, as indicated on the plot of vesicle diameters for this class (**I**, circled in red). See video S5. Scale bar: 500 nm **(A)**.

**Figure 6.**
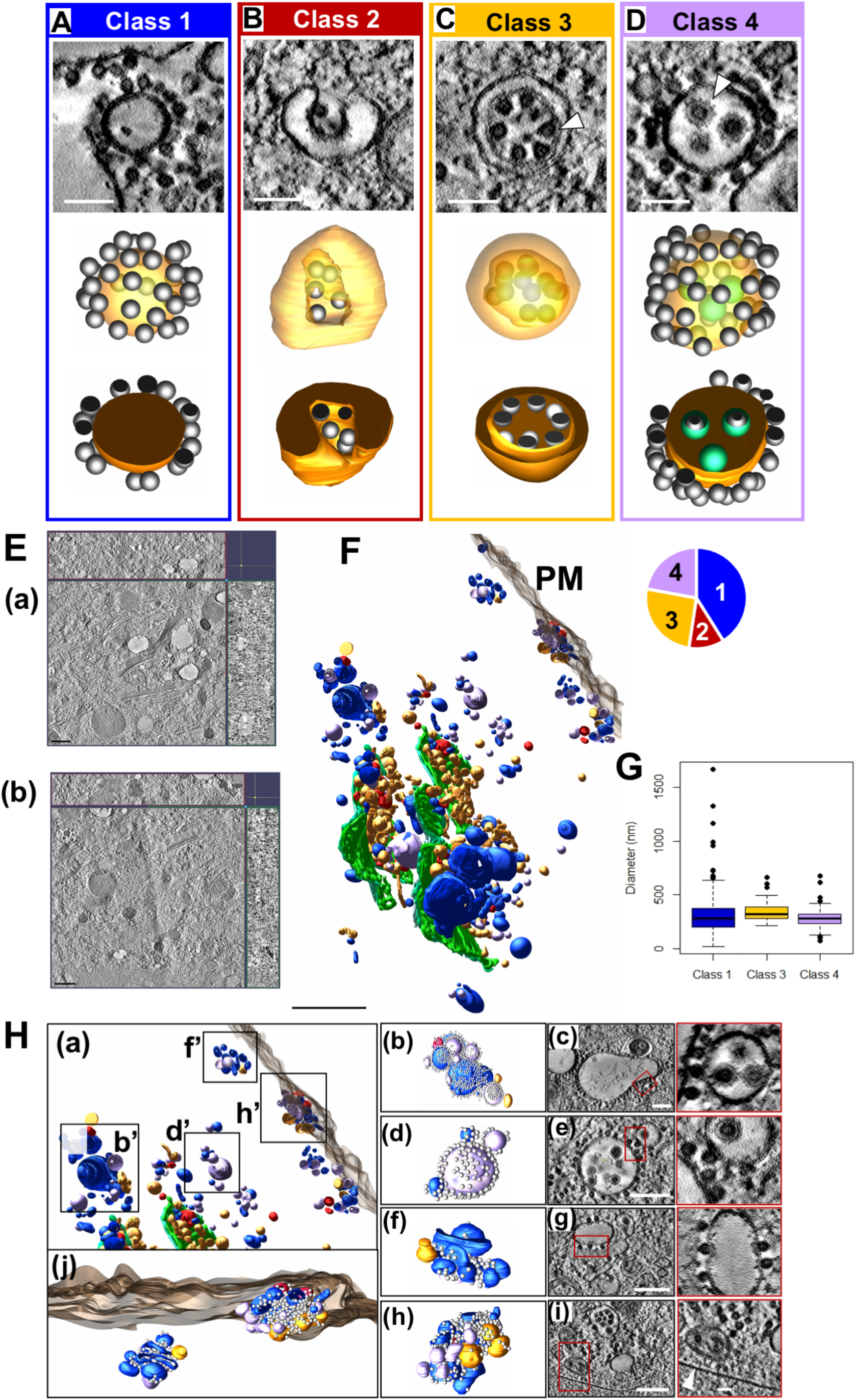
Classification and three-dimensional analysis of four morphological forms of CPV-II. **(A-D)** A representative vesicle from each of the four CPV-II classes observed in the large volume tomogram are shown in three views: 3-nm thick virtual sections (top image), full 3D models made partially transparent (middle image), and 3D cutaway models (bottom image, See **videos 6-9)**. NC **(C, arrowhead)** is distinguishable from fully enveloped virus particles **(D, arrowhead)**. A 3-nm thick virtual section from tomogram 1 and tomogram 2 **(E**, (a) and (b) respectively) with z-depth along the x- and y-axis shown along the top and right side, respectively that were joined together for analysis of CPV-II distribution (See also Fig S7). **(F)** Spatial distribution of segmented CPV-II forms in the combined tomographic volume are shown in reference to the *trans*-most Golgi cisternae (green) and the plasma membrane (translucent grey). CPV-II are color-coded by class: class 1 (blue), class 2 (red), class 3 (gold) and class 4 (lavender) (see video S10). The pie chart in the upper left shows the fraction of vesicles in each class. **(G)** For the three classes of roughly spherical CPV-II, the volume of each vesicle was calculated from the segmented mesh, and a diameter was approximated for each vesicle. Size distributions for class 1, 3, and 4 CPV-II are represented as a box plot with outliers representing data points beyond 1.5 times the interquartile range. Scale bars: 100 nm **(A-D)** and 1 μm **(E). (H)** CPV-II clusters in the cytoplasm as well as at the plasma membrane indicated by b’, d’, f’, and h’ in panel (a). Several vesicles in the cluster (h-i) come in proximity to the plasma membrane [white arrowhead (i)]. Magnification of the region outlined in red (c), (e), (g), and (i) depict the interface between vesicles within the cluster. In C and G, multiple NC binds two separate CPV-II membranes through bivalent interactions; an observation that explains the clustering of CPV-II. Scale bars: 200 nm, (c, e, g, i).

**Figure 7.**
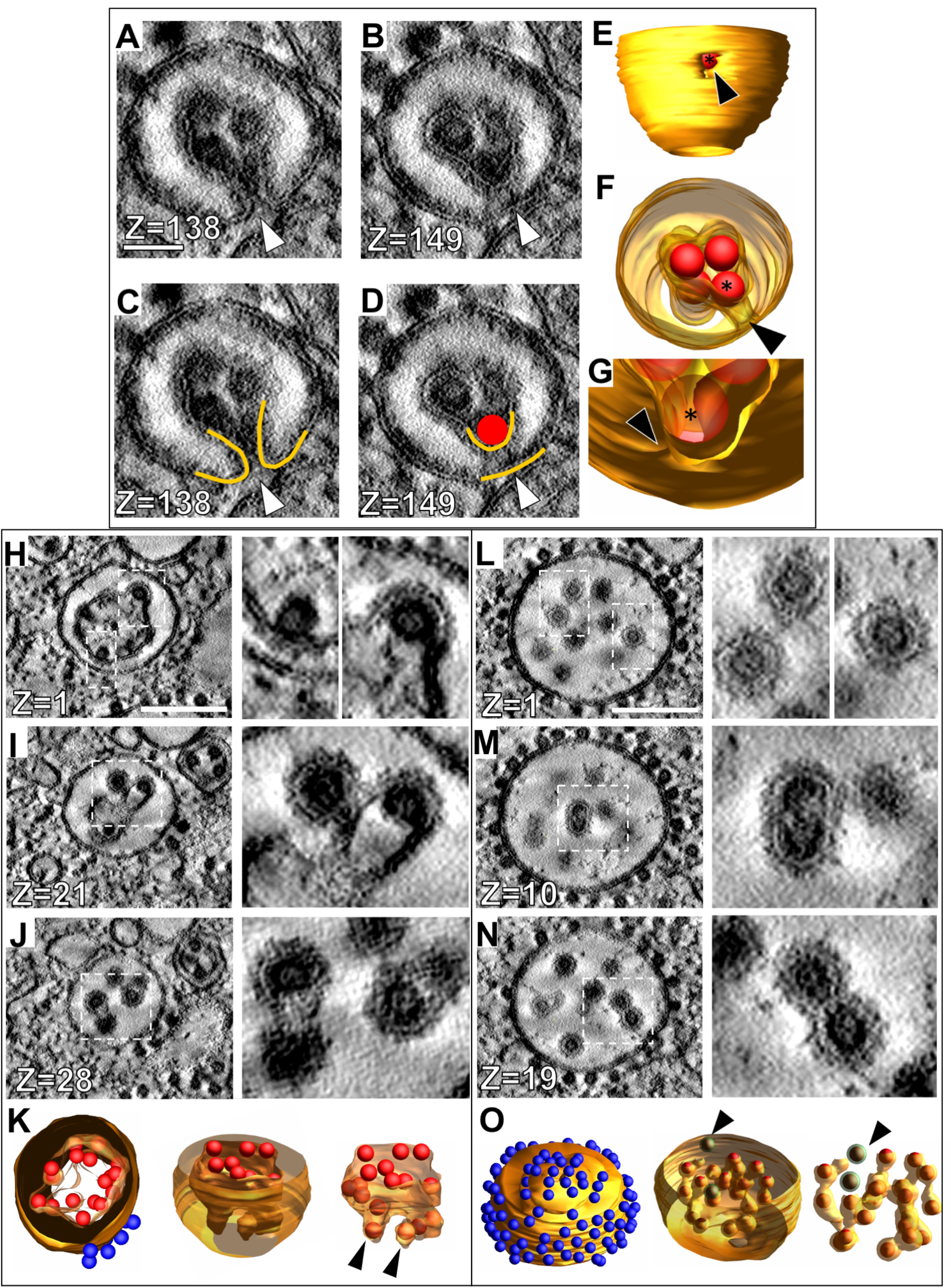
3D analysis of intermediate forms between classes 2, 3, and 4 reveals a potential maturation pathway. **(A-B)** Tomographic slices through an intermediate between class 2 and class 3 CPV-II. Each z-slice is 3 nm in thickness, with the slice number shown in the lower left corner of each image. The fusion pore is indicated by the white arrowhead in A and C. **(C-D)** Tracing of the vesicle membranes (gold) in A-B and a NC (red sphere) near the fusion pore. **(E-G)** 3D rendering of the vesicle shown in A-D with the fusion pore indicated by a black arrowhead. See Video S11. **(H-K)** Tomographic slices at various z-heights of a CPV-II which is intermediate between class 3 and class 4, showing various stages of interior NC envelopment, from moderate amounts of budding **(H)** to nearly complete envelopment **(I-J)**. White dotted lines delineate areas magnified to the right. **(K)** 3D rendering of the vesicle shown in H-J. See Video S12. **(L-O)** Tomographic slices through a class 4 CPV-II containing enveloped virus particles **(L)**, dual-core particles **(M)**, and enveloped NC connected to each other by membrane **(N). (O)** 3D rendering of the vesicle shown in L-N, illustrating the interconnected nature of the partially enveloped NC. Completely enveloped NC without any membranous connection is indicated by the green segmentation and black arrowhead. See also video S13. **Scale bars:** 50 nm **(A)** and 200 nm **(H, L)**.

**Figure 8.**
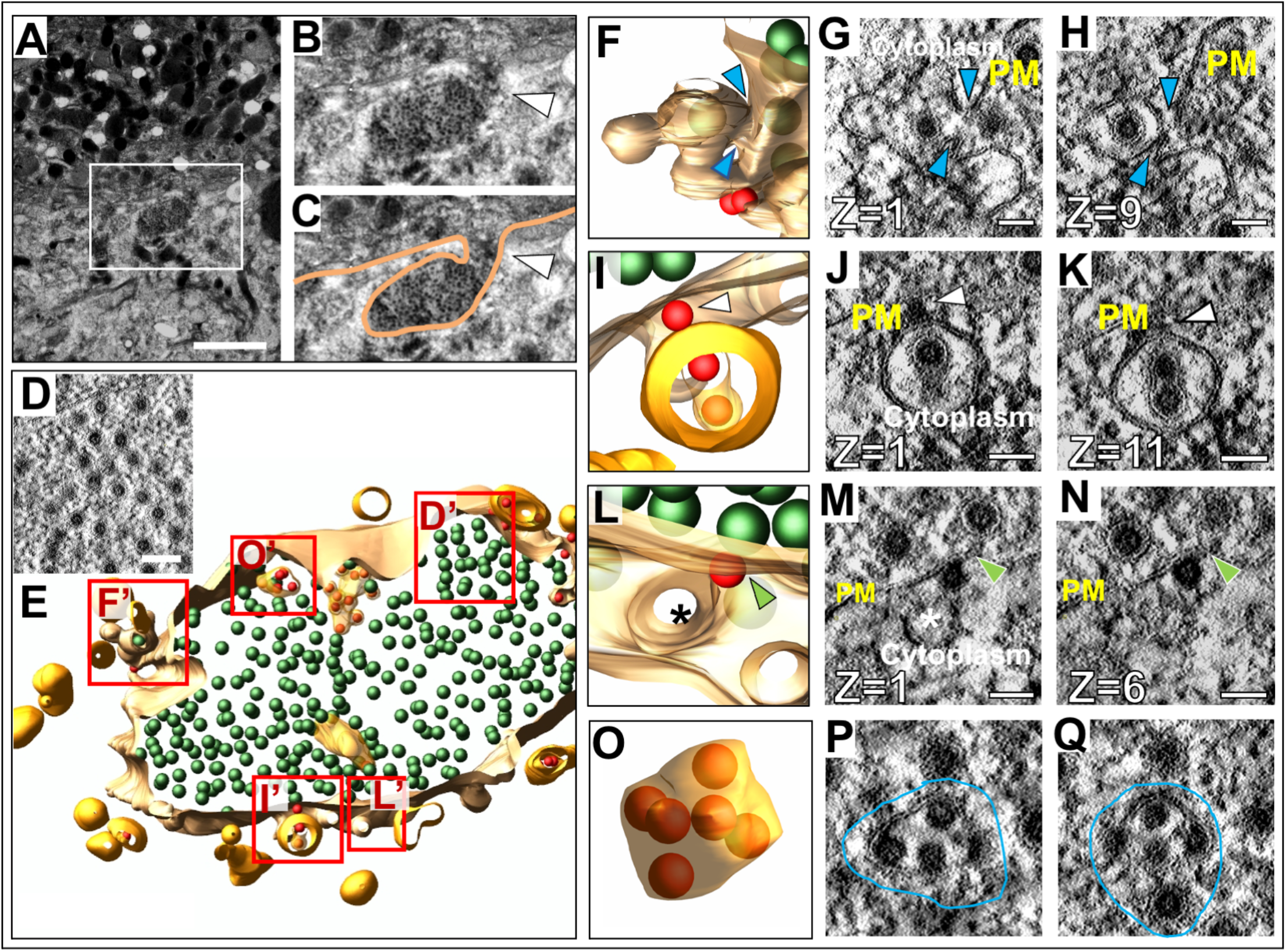
Interaction of CPV-II with the inner leaflet of the plasma membrane. **(A)** TEM image of an area of a large vesicular structure connected with the plasma membrane via a narrow neck/tunnel like extension. **(B)** Magnified area of the yellow-dotted outline in **(A)**. White arrowhead indicates the opening to the extracellular space. **(C)** Tracing of the plasma membrane shown in **(B). (D)** Tomographic slice through the sac-like structure shown in **(A-C)**, revealing that it is filled with virus particles. **(E)** Segmented model of a 250-nm thick slice of the sac-like structure, with areas labeled D’-O’ magnified in panels **(F-H)**. 3D rendering and slices through different z-heights of a class-4 CPV-II fusing to the plasma membrane. Blue arrowheads indicate the continuity between the vesicle membrane and the plasma membrane indicating a neck-like opening at a different z-heights **(I-K)**; another class-4 CPV-II bound to the plasma membrane via a surface NC. White arrowheads indicate the NC simultaneously binding the plasma membrane and the CPV-II membrane, Video S14. **(L-N)** A Class 1 CPV-II with a NC in the initial stages of budding through the plasma membrane. Green arrowheads in M and N indicate the bending of the inner leaflet of the PM at the NC binding site (See Video S15) **(O-Q)** An extracellular vesicle containing incompletely budded NCs in various stages of envelopment (outlined in blue) found within the sac-like invagination. Scale bars: **(A)** 2 μm, **(D)** 100 nm, **(G-Q)** 50 nm.

### Outward bending of Golgi cisternae that bind viral NC provide clues for the biogenesis of double lamellar CPV-II

The most prominent feature of VEEV induced Golgi remodeling as seen in conventional 2D TEM studies is the herniation of cisternae and formation of large vacuoles (∼0.2-1.5 microns) that perhaps in time separates out to form single lamellar CPV-IIs (Fig S2). However, these studies provide little information on the morphogenesis of other classes of CPV-II that were identified in our Golgi marker study (Fig 1). Thus, we looked for subtle structural clues on the segmented tomogram of the Golgi to address this. The two Golgi stacks under study (stack-1 and stack-2, Fig 3A, B, E and F) both show the presence of viral NC on the first intact cisternal face. Many of these NCs were bound to the cisternal rims (edges) as illustrated for stack-1 and stack2. The edges displayed curvature away from the plane of the stack at the locations where the NCs were bound (Fig 3A-H, Video S2). Close examination of the bent edges of the cisternae revealed that the section that bent around NCs largely retained the narrow intralumenal distance of cisternae present in a Golgi stack (Fig 3I-L). A large curvature of the cisternal rims was composed of multiple bending events occurring around bound NC (Fig 3M-P). However, not all bound NC were associated with cisternal bending. This was true especially with the ones in the middle of the stack. In order to visually clarify membrane bending, we chose to highlight any bending that was possibly associated with cisternal wrapping around the NC by selecting NCs located on membrane which had a local radius of curvature between 20 and 60 nm, corresponding to the 40 nm radius of a NC (Fig S5). These NCs were colored red in the 3D model. NCs located on membranes with radius of curvature greater than 60 nm were colored blue.

In addition to curling of the cisternal rims, a single large cup-like transformation of a Golgi cisterna was observed (Fig 3Q-S and Video S3). This formation consisted of a largely intact cisternal structure curving away from the plane of the Golgi stack (in light green, Fig 3Q and R), with one edge closing to form a vesicle-like structure (top-down view in Fig 3S). This form essentially represents a transitional intermediate between a flat Golgi cisterna and a double membrane vesicle. A high concentration of NCs is present on the inner membrane of this intermediate, but only 14 out of 97 NCs show membrane curvature indicative of a potential membrane wrapping event (red spheres in Fig 3S). This intermediate largely retains the characteristic intralumenal distance of a Golgi cisterna (15-19 nm) and provides the most telling evidence for the morphogenesis of double lamellar CPV-II.

### Pleomorphic membrane structures with bound NC are fragmented Golgi cisternae

Immediately adjacent to the *trans-*face of stack1, large pleomorphic membrane bound structures were observed (segmented tomogram in Fig 4A, Video S4). The structures closest to the Golgi stacks appeared to be discrete structures in single slices through the tomographic volume (Fig 4B, pink and brown), but 3D segmentation revealed that they are in fact partially intact Golgi cisternae still somewhat aligned with the stacked cisternae (Fig 4C-D). The central part of the structure shown in Fig 4C (area containing bound blue NC) retained flat cisternal morphology and maintained contact with the preceding cisterna in the stack (Fig 4B, arrowhead), whereas the outer edges exhibited a fenestrated morphology (Fig 4C arrowheads). Two small double-lamellar vesicular structures surrounding small groupings of NC were also present (Fig 4C, inset). The next closest cisternal structure, in brown, exhibited more vesicular structure than the previous, with long tubular elements connecting the vesicular formations (Fig 4D).

Moving further away from the *trans*-face, pleomorphic CPV-II structures were observed to contain a combination of small vesicular formations and tubules with highly varied morphology (Fig 4E-H). Some of these structures included up to ten connected vesicular formations, resulting in structures resembling grape bunches (Fig 4E-F). Other structures consisted of a few vesicles with tubular connections (Fig 4G), or even simpler; a single vesicle with a tubular component (Fig 4H). Tomography proved to be the key to unraveling their complex morphologies and connections (see corresponding 2D slices in Fig 4E-H).

### Three-dimensional analysis of Golgi stacks associated with large unilamellar CPV-II

One of the Golgi stacks had closely associated large vesicles, reminiscent of those observed at earlier time points by 2D EM and shown to be of GA origin (Fig 5A, compare with Fig S3A(v) and B(vi). While the presence of NC on the vesicles in 2D was either not observed (at early time points) or perhaps went undetected, the large vesicles in the tomogram clearly have large numbers of NC bound to their cytoplasmic surfaces, indicating that they are indeed CPV-II. Particularly, a large irregularly shaped vesicle of approximately 1.5 µm diameter in close apposition with the *trans*-most Golgi cisterna of the stack (Fig 5B and C). This vesicle was separated from the cisterna by a single layer of NC which is bound to both the CPV-II and the Golgi membrane in a bivalent association (Fig 5C-E). Segmentation and 3D visualization of this vesicle and cisterna revealed increased NC density in the region where the vesicle was bound (Fig 5F-G).

The visible deformation on the Golgi cisterna where the CPV-II was positioned further confirmed the tight association of the CPV-II with the Golgi (Fig 5G). Examination of the region immediately surrounding these large vesicles in contact with the Golgi, revealed several additional large, irregularly shaped CPV-II, as well as closely associated smaller CPV-II (Fig 5H and Video 5). Vesicles were apparently connected into a large cluster by NC bivalently bound to two vesicles (Fig 5C black arrowheads). While most of the vesicles in the cluster fell within the typical size range of CPV-IIs, it is noteworthy that the three large vesicles in this cluster were the three largest CPV-IIs found in the whole tomographic volume, but distinct outliers in terms of size (Fig 5I, circled data points). The vesicle highlighted in Fig 5F and found in closest apposition with the GA is the largest CPV-II of the 353 analyzed in this volume. From our observations from the marker study as well as these rare large CPV-II in the tomogram, it is tempting to interpret these large CPV-II as portions of herniated Golgi cisternae. It is unknown whether these large irregularly shaped vesicles persist or are unstable intermediates that may break down into the smaller, more spherical shaped class I CPV-II observed throughout the volume.

### Identification and three-dimensional analysis of four morphological forms of CPV-II

To date, published ultrastructural data on CPV-II has been obtained via traditional thin section studies. Due to the complex morphological nature of CPV-II it is difficult to ascertain its real 3D form by thin section screening. In addition, since Golgi are an architecturally complex organelle, we undertook a comprehensive 3D study on CPV-IIs utilizing 3D data from two overlapping tomograms to investigate the morphological pathway of the origin of CPV-IIs (Fig 6E a and b). As previously mentioned, a CPV-II is defined as alphavirus induced vesicles with NCs bound to its membrane that predominate during mid to late-stage infection in mammalian cells. In these tomograms, we observed 353 discrete vesicles that can be categorized as CPV-II. After segmentation and 3D visualization of each CPV-II, we identified four distinct morphological forms of CPV-II. First, class 1 CPV-II were defined as unilamellar vesicles with NC bound to their cytoplasmic surface (Fig 6A and Video S6). Class 2 was composed of vesicles containing an invagination with NC bound to the membrane in the invaginated space (Fig 6B and Video S7). Class 3 contained bi-lamellar vesicles, with NC bound to the inner membrane and variably present on the outer membrane (Fig 6C and Video S8). Finally, class 4 vesicles were defined as single-membrane vesicles with enveloped virus particles inside and NC variably present on the outer membrane (Fig 6D and Video S9).

### Intracellular distribution of CPV-II

To investigate if the subcellular distribution of the four CPV-II classes follows a specific pattern they were next analyzed for spatial distribution, size and distance from the nearest Golgi stack. In order to increase the sampling size of each of these 4 classes and to understand their distribution a second tomogram (Fig 6E b) was collected that overlapped with the first tomogram (Fig 6E a) together extending volume up to the plasma membrane (Fig S4). 3D data on the CPV-II types was then collected from the joined segmented tomogram (Fig S4, D). Each CPV-II was then color-coded by its morphological class and shown with the *trans*-most Golgi cisterna of the relevant Golgi stacks and the PM for spatial reference (see also Fig 6F) and Movie S10). Of the 353 CPV-II, class 1 was the most abundant at 145 vesicles (41%), followed by class 3 (90, 26%), class 4 (78, 22%) and class 2 (40, 11%) (Pie-chart in Fig 6F). The four classes of CPV-II were also analyzed for distance from the nearest Golgi stack to determine the occurrence of these classes in CPV-II clusters proximal to the Golgi and at the plasma membrane (Fig S6, E).

The sizes of classes 1, 3, and 4 were also analyzed by calculating the volume of the segmented volumes for each vesicle using IMOD (Kremer et al, 1996), and converting volume measurements to diameter assuming a spherical shape (Fig 6G). For class 3, the outer membrane was used for size calculations. CPV-II in classes 1, 3, and 4 had similar diameter distributions for the interquartile range, with means of 326, 340, and 286 nm, respectively. Class 1 exhibited a notably broad size distribution with diameters ranging from 22 to 1667 nm, although a few extremely large vesicles were observed. Class 3 had no vesicles smaller than 214 nm, and a narrow size range up to 662 nm.

CPV-II are distributed all over the cytoplasm including the inner leaflet of the PM. Interestingly, CPV-II exists primarily as clusters, especially at the plasma membrane. These clusters were shown previously to carry the Golgi marker alpha mannosidase-II ratifying their Golgi origin (Fig 1B-C). A careful analysis of the clusters revealed that irrespective of its location, surface NC within these clusters made bivalent contacts between two adjoining CPV-II possibly holding the cluster together. Two representative clusters closer to the Golgi (Fig 6 (a)b’, (b) and (c), and d’, (d) and (e), respectively) and two by the PM (Fig 6H (f’), (f) and (g), and h’, (h) and (i), respectively) are shown. CPV-II clusters at the plasma membrane (depicted in Fig 6H(a) are shown as a segmented model from different point of view (Fig 6H(j), see Video S10)

### 3D analysis of intermediate CPV-II forms reveals a potential maturation pathway

In addition to the 4 classes of CPV-II just described, forms exhibiting intermediate features between the defined CPV-II classes were also observed. We therefore examined these forms closely for obtaining clues about their relatedness. First, the forms exhibiting characteristics of both class 2 and class 3 were examined. A representative vesicle depicted in Fig 7A-G (and Video S11), containing small areas of horse-shoe shaped cross-sections like class 2 (Fig 7A and C, compare with Fig 6B), while most of the vesicle cross-section resembled class 3 (Fig 7B and D, compare with Fig 6C). 3D visualization of the vesicle clarified the largely double-lamellar nature of the vesicle, apart from a small pore-like opening from the inner NC-containing compartment to the extravesicular space (Fig 7E arrowhead). The opening consisted of a narrow channel (Fig 7F-G arrowheads) with a single NC present inside the opening (asterisk in Fig 7E-G).

Apparent intermediates between class 3 and class 4 CPV-II carrying partial characteristic of both were also observed and analyzed in 3D. Tomographic slices through one such vesicle revealed slices resembling class 3 bi-lamellar vesicles (Fig 7H), other slices with nearly enveloped virus particles resembling class 4 CPV-II (Fig 7J), and intermediate slices with NC which appear to be budding into the inner membrane of the vesicle (Fig 7I). 3D visualization revealed a single inner compartment with many of the NC in various stages of apparent envelopment, but all still connected to the main compartment (Fig 7K, arrowheads, Video S12). The wrapping/envelopment of NC by the inner membrane of class 3 CPV-IIs became very apparent in 3D and this remodeling of the inner membrane around the NC trapped within was a common observation in the members of Class 3. This observation suggested that the double lamellar structure may not be the final end-product of this pathway and that further remodeling of the inner membrane possibly occurs.

A second intermediate between class 3 and 4 is shown in Fig7 L-O (see also Video S13). In the 2D tomogram slices, this vesicle that appears to be a class 4 vesicle with mature enveloped virions within; however, reveal membrane connections between nearly all the membrane wrapped NCs, indicating that they are not mature virions as concluded initially from thin section screening (Fig 7M, N). Segmented tomograms clearly show the membranous connections between almost all the wrapped NC except for two (Fig 7M-N, NC in green shown with an arrowhead in O) show lack of detected membrane connection with the rest of the membrane wrapped virions)

To determine if CPV-II clusters play a role in virus egress we screened the cell periphery to visualize CPV-II fusing with plasma membrane that would suggest that. In the process we came across large (∼1 × 1.5 μm) sac-like structure located at the PM filled with a high density of virions within (Fig 8A-E). This sac-like structure revealed a continuity with the plasma membrane and an opening to extracellular space via a narrow ∼700 nm long neck (Fig 8B and C, white arrowhead). Various forms of CPV-II were seen closely interacting with the inner cytoplasmic leaflet membrane of this structure (Fig 8E). Careful scanning of the different Z slices revealed a class-4 CPV-II in what appeared to be in a fused state with the PM (Fig 8 F-H and F’ in E, see Videos S14-S15). A second class-4 CPV-II with a dual-cored virus particle was seen bound to the PM via an external NC (segmented in Fig 8I and its tomographic slices J and K). A class-1 CPV-II was also observed bound by an external NC to the PM (Segmented in Fig 8L and in its tomographic slices M and N, respectively). The PM at the site of interaction with the NC exhibits bending, possibly indicating an initial stage of budding (Fig 8M and N, green arrowheads). Despite these observations, fusion-like events were rare (<10) in occurrence in tomogram 2. We next examined the contents within a large sac-like structure to determine to study its content. Segmentation revealed mostly fully formed free virions within the sac. Interestingly, in addition to virions several single lamellar vesicles containing multiple NCs in various stages of envelopment were also observed that were reminiscent of incompletely enveloped NC within the lumen of the Class-4 CPV-II (compare Fig 8 O-Q with Fig 7L-O).

## DISCUSSION

In this work, we sought to answer the question of how does stacked cisternal architecture of the Golgi get transformed into the vesicular forms CPV-II during alpha virus egress; a question analogous to the ones asked about a host of similar processes such as organelle biogenesis and morphological transformation seen of organelles in various diseases. This structural remodeling of secretory organelles of cells infected with enveloped virus overlaps temporally and mechanistically to its egress pathway (Risco et al, 2003; Salaneuva et al, 2003; Welsch et al, 2007, Welsch et al, 2009;; Johnson et al, 2011). The viral components and interacting host proteins linked directly to its egress pathway has been the focus of intense research for decades, however ultrastructural studies revealing the structural manipulation of the secretory organelles is relatively scarce. Such studies are critical for understanding the viral subversion of the host cell secretory system during egress as well as clues for structure-function relationship of these highly structured organelles.

This required imaging of large cellular volumes at a resolution sufficient to detect perturbations in the lipid bilayer. In order to reach such a goal, we understood that the flexibility of repeated data collection from the same sample/cell is the key. Thus, resin based electron tomography (ET) of cryofixed sample was pursued and optimized (Ladinsky et al, 1999; Ladinsky et al, 1994; Miranda et al, 2015; Perkins et al, 1997). A similar approach (ET of serial-sections or ssET) was previously employed to image up to an entire at a resolution of about 15 nm (Marsh et al, 2001; Noske et al, 2008). Here we implemented an improvised phased data collection from 250nm resin serial-sections and utilized the montaging function of the serial EM software cover a larger area at a targeted magnification. Traditional EM imaging and Golgi-marker localization indicated the Golgi origin of the morphological forms of CPV-II. However, we were mindful that since GA, especially the trans-Golgi is a major sorting junction where vesicles from the PM and the endosomes constantly communicates making it an entity with a mosaic of membranes of different origins. Thus, although the structural genesis of CPV-II may begin at GA, it is expected contain some membrane components and even markers from organelle like the endosome, ER etc. This is especially true if the GA in its CPV-II avatar still retains its canonical functions. As a result, we have kept this paper limited to dissect the contribution of the Golgi apparatus in the biogenesis of CPV-II. A more comprehensive 3D marker study is ongoing that aims to dissect the contribution of other organelles in the biogenesis of CPV-II.

Prior work in the field has employed traditional thin-section EM and was able to identify the unique NC-studded CPV-II structure (Grimley et al, 1968; Mussgay & Weibel, 1962; Soonsawad et al, 2010). However, the information obtained from these studies by the discrete and limited sampling of membrane could not establish the direct relationship of the cytopathic vesicles (CPV-II) with the GA. Thus, to detect these putative early changes and to determine the relationship of CPV-II with the Golgi apparatus, we carried out HRP-tagged marker studies employing a recently published method for superior sample preservation during localization of peroxidase tagged proteins (Sengupta et al, 2019). Using HeLa cells constitutively expressing HRP-ManII we demonstrated that this Golgi perturbation during alphavirus infection is neither cell type specific, nor an artefact resulting from overexpression of HRP-ManII. However, two things were apparent from our thin section 2D analysis; first, the GA begins to lose its cisternal architecture via widespread herniations and vacuolization leading to the formation of large pleomorphic vesicles as early as 3 hours PI. Second, thicker serial sections showed that CPV-II form large clusters that could have extensive connections with each other and may not all be isolated vesicles as previously thought (Grimley et al, 1968; Mussgay & Weibel, 1962).

We carefully analyzed multiple examples of each of these subclasses to look for intermediate forms that showed characteristics of two or more classes. The first tomogram covered a large subcellular volume consisting of multiple Golgi stacks where *en bloc* remodeling of cisterna was apparent. In order to record these changes systematically, our analysis progressed outward step by step from *trans-*most intact cisternae. The predominant CPV-II observed in our thin section (Fig S2), Golgi marker study (Fig 1) as well in our tomograms (Fig 6) possibly originate from vacuolized Golgi cisterna as described previously by Griffith*s et al*. The other type of Golgi remodeling distinct from the former, results in CPV-II class 2, 3 and 4. We provide 3D ultrastructural evidence that this second type originates from the bending and curling of intact Golgi cisternae. Transitional intermediates of various sizes exhibiting part cisternal and part bi-lamellar vesicular characteristics on the Golgi further substantiates this initial phase of the proposed morphological pathway (Fig 3Q-S and 4E-H).

Further away from the GA, independent CPV-II was analyzed in 3D for putative links. Intriguingly, many of these vesicles that at first appeared to be Class 2, 3 and 4 CPV-II were found to exhibit characteristics of two classes simultaneously. The ultrastructural evidence points to Class 2 as the precursor of class 3 (Fig 7). These intermediates seem to have been captured at a stage just prior to end-to-end fusion giving rise to *bona fide* class 3 double lamellar CPV-II. This characteristic of bent Golgi cisternae to form bi-lamellar structures by end to end fusion to create a compartment within a compartment has also been recently reported in brain tissue, where these Golgi structures function as a degradosomes (Fernandez-Fernandez et al, 2016).

The morphological relationship between class 3 and class 4 may be the most intriguing of all. Fig 9H-K shows a typical example of such a transitional CPV-II that were in some of the 2D slices. This resembles class 3 CPV-II where the NC are just bound to the inner membrane without any visible distortion of the membrane or budding. However, advancing through the z-slices for the same CPV-II, it becomes apparent that the inner membrane at the point of contact with the NC exhibits bending consistent with various stages of alphavirus NC envelopment. Video S12 gives a great example of this feature. This observation led us to carefully examine the lumenal content of Class 4 that apparently contains fully enveloped virions (Fig 7L-O). To our surprise, we observed that only two of the enveloped NC within this CPV-II are independent virions. The rest of the seemingly matured virions were found to be connected to each other via intact membrane connections. This indicated that either their envelopment was incomplete at the time of cryofixation, or they never mature to form independent virions and that so far these membrane connections have gone undetected. Thus, our model proposes that Classes 2, 3 and 4 CPV-II are all intermediates of the same morphological pathway that gives rise to enveloped/membrane wrapped virions during alphavirus infection in mammalian cells (Fig 9).

**Figure 9.**
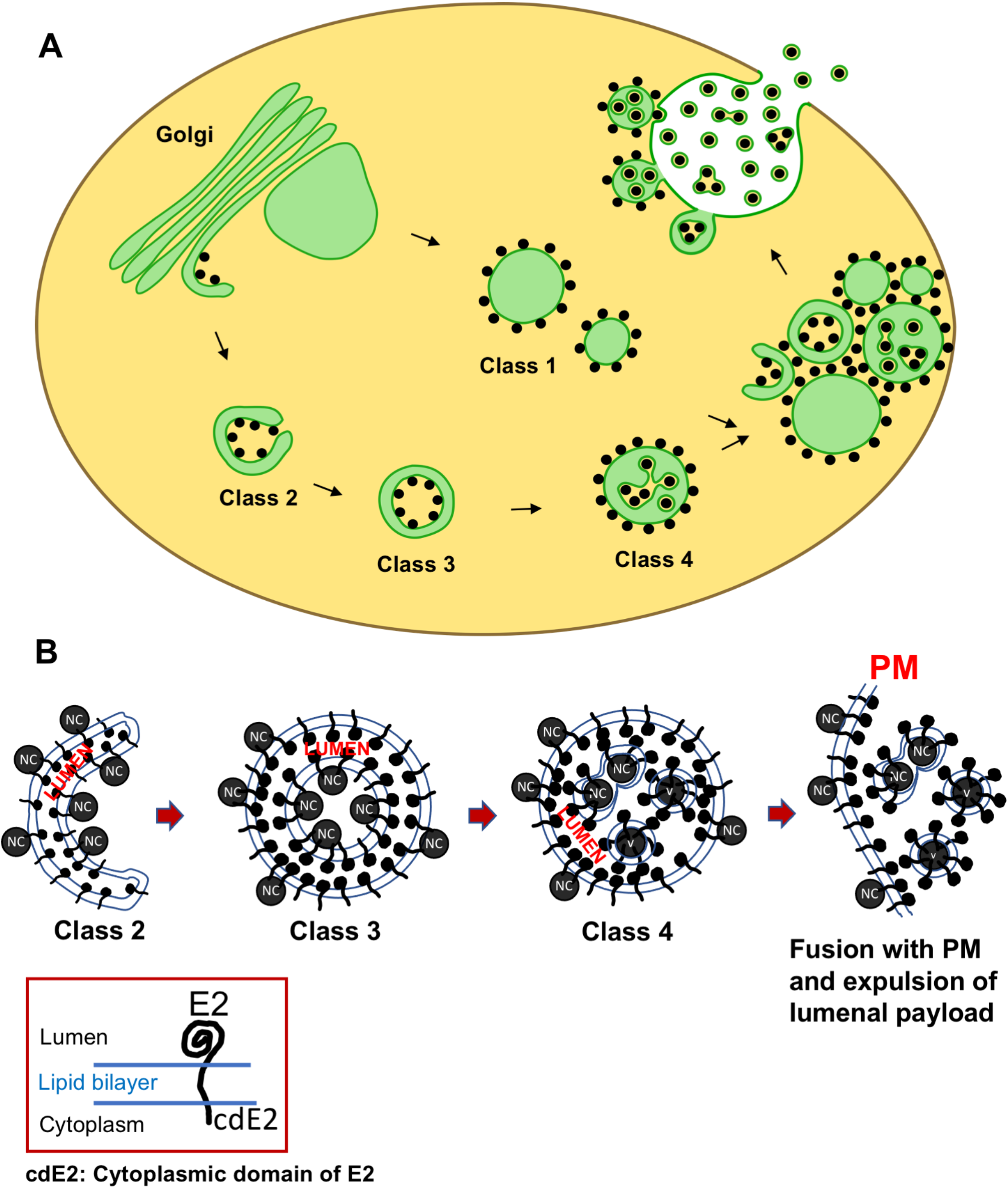
Model of Golgi remodeling resulting in the biogenesis and maturation of CPV-II. **(A)** Proposed model illustrates that despite the distinct presence of four morphological classes of CPV-II; potentially two mechanisms operate. The morphogenesis of class 2, 3 and 4 begins with the Golgi cisternae bending and fragmentation. Class 1 CPV-II, on the other hand, forms directly by herniation and swelling of the Golgi cisterna. Class 2 (or the sickle form) bends onto itself and fuses to give rise to the double lamellar structure (DMV) of Class 3. Finally, NC trapped within the DMV acquires its envelope from the inner membrane of the DMV to result in class 4. Finally, all these forms clusters via bivalent interaction of surface NC that bridges numerous CPV-II together and are trafficked to the plasma membrane. From our data on events at the plasma membrane, we speculate that the NC on the surfaces of CPV-II is first transferred to the inner leaflet of the PM from where they bud out. In parallel, the CPV-IIs much smaller in size, fuses and delivers the lumenal virions to the cell exterior. **(B)** Hypothesizes the orientation of the E2 glycoprotein in the different classes of CPVII as well as the presence of NC and their potential delivery to the plasma membrane.

To lend more credibility to our hypothesis of inter-class conversion and maturation of CPV-II, we calculated the frequency of each class as a function of their distance from the Golgi (Fig S6E). It was observed that the largest class, Class 1 remained the predominant class and its counts comparable irrespective of its distance from the GA. However, as the distance from GA increases, the class 2 and 3 CPV-II counts fell considerably while the count for class 4 increased. This observation supports our class conversion hypothesis in two ways. One, that class 2 and 3 are possibly intermediate forms that mature to class 4 as they are trafficked towards the PM in clusters, and two, the morphogenesis of Class 1 occurs by a separate mechanism and not by bending of the Golgi cisterna. Indeed, our results from the thin section screening, Golgi marker localization, as well our tomograms provide evidence that Class 1 could in fact form from swelling, vacuolization and fragmentation of the Golgi cisterna observed from a very early stage of infection (Fig S2(iii-iv), 5(A, B and H), S2(1-4) and S4(iv-v).

One very apparent feature of the CPV-II analyzed within these overlapping volumes, is that they exist as large clusters, more so at the plasma membrane (Fig 6, H). CPV-II within these clusters are apparently tethered via bivalent interaction of NC (Fig 6H(b-i)). Thus, a series of such interactions between multiple core-carrying vesicles could result in the formation of an elaborate vesicular cluster that can traffic as one. This mode of trafficking would be advantageous to the virus as it serves as an efficient mechanism for delivering large reservoirs of both viral glycoprotein and NCs from the GA to localized sites on the plasma membrane creating micro-domains rich in viral structural protein to facilitate plasma membrane budding. Recent evidence demonstrating the requirement of actin remodeling proteins ARP3 and RAC1 in the later stages of infection in addition to results showing the direct interaction between actin and E2 using biochemistry and electron microscopy, indicate that CPV-II trafficking to the PM is actin dependent (Radoshitzky et al, 2016)

To gain a quantitative insight in this putative intracellular envelopment in alphaviruses, we investigated the potential efficiency of virus envelopment via this process. To this end, the surface areas of the inner compartment of Class 3 CPV-II were measured and the number of NCs inside were counted to determine if there is enough membrane for the internal NCs to become fully enveloped. We approximated that 5000 nm^2^ (dotted line, Fig S5A) of membrane is required to envelope each NC (approximation based on a 40 nm diameter of NC). For class 3 CPV-II, we plotted the lumenal membrane surface area available per NC versus the percent of NC that were undergoing potential budding. For each NC bound to the inner membrane, budding was determined as before based on the radius of curvature of the membrane at the point of attachment (Fig S6). In general, the more membrane available, the more NC exhibited budding; however, 53% of the CPV-II (9/17) did not have sufficient membrane to envelope all internal NCs (Fig. S7). Thus, it is estimated that half of all NCs inside double lamellar CPV-II could not become fully enveloped virus. This may point to the fact that this could be a dead-end byproduct of the formation of bi-lamellar CPV-II.

Several intriguing questions emerge from our results. From the virology standpoint the primary question, is this pathogenic conversion of Golgi has any bearing with the virus egress pathway? The answer obtained through this study is not clear cut. Our findings indicate that exocytic expulsion of internally budded virions is probably not a very efficient process judging by the number of Class 4 CPV-II observed in the state of fusion (<10) with the PM. Interestingly a higher frequency of NC on the outer surface of CPV-II (Class 1 to Class 4) are observed making direct contact with the PM and bending the inner leaflet of the PM (Fig 8(I-N) and represented in the model, Fig 9). One way to explain these observations is that the CPV-II can potentially mediate two pathways of viral egress at this stage of infection. First, the NC on its surface could potentially be transferred to the inner leaflet of the PM inducing plasma membrane budding and second, this initial interaction of the CPV-II via the NC on the surface could in turn facilitate the fusion of the CPV-II leading to its fusion to the PM and expulsion of the lumenal virus payload. In summary, despite the apparent inefficiency of the exocytic pathway, the CPV-II cluster that carries much greater number of NC on its surface than in its lumen, brings in a massive cloud of NC and envelope glycoprotein to the PM that at least feeds into the major budding pathway via the PM. Lastly, is also possible that the low number of budding/fusion events detected at the CPV-II clusters could be a result of the sample preparation. In this regard, cryoEM methods are being pursued currently to carefully screen the plasma membrane at the same stage of infection to rule out the possibilities.

As tools for visualizing virus induced membrane modifications *in situ* gets more and more sophisticated more examples of viruses with multiple modes of budding and egress is gradually coming to light (Saraste et al, 2021; Ghosh et al 2020, Spiropoulou et al, 2001). In several virus families such as alphaviruses, whether a virus buds into intracellular membrane or at the PM is dictated largely by the interaction between the cytoplasmic domain of the viral glycoprotein (spike) and the cytoplasmic NC. For example, in alphaviruses chemically induced accumulation of the glycoprotein at the Golgi results in the binding of NC directly to the Golgi and budding-like events (Griffiths et al, 1993). Thus, in our study the late stage in VEEV infection could represent a stage where natural accumulation of glycoprotein at the Golgi results in the binding of NC and formation of CPV-II. This in turn could be the mechanism by which the virus induces a more efficient co-trafficking of glycoproteins and NC to the PM in these late stages. In this regard, the CPV-II cluster may not be directly fusing to the PM could rather be seen as sort of virus induced “Golgi outpost” that could make the transfer of NC and glycoprotein more efficient as more and more viral proteins builds up within the host cell. Further sorting and budding possibly operates within these clusters. Two observations support this theory; one, the absence ManII-HRP signal on the PM in our DAB based localization indicates the lack of/inefficient fusion of large CPV-II clusters that we show carrying the enzyme and two, absence direct visualization of multiple fused CPV-II with the PM despite the presence of multiple clusters. Additionally, the CPV-II involved in the rather infrequent fusion events (<10) with the PM are around 100nm in diameter indicating that a further maturation and sorting state could be operating from these CPV-II clusters. In this context it will be interesting to investigate further whether the CPV-II that are technically remodeled Golgi cisterna still retain its secretory functions. Remodeling of the GA to create secretory Golgi outposts as result of changes in secretory demands of cells has been reported (Hanus et al, 2008; Kreft et al, 2010; Višnjar et al, 2017). Thus, mobilized Golgi cisternae in the form of CPV-II that clusters near the PM could be an example of viral subversion of this program to induce secretory outposts that function to meet the elevated level of structural protein transport to PM during budding. Either way, the study of Golgi remodeling in viral infection is also critical in understanding viral pathogenesis. Suppressive effect on the endogenous secretory system could lead to the inhibition of inflammatory cytokines, anti-viral interferons and cell surface expression of MHC-1. Remodeling and inhibition of the secretory pathway has been cited as a mechanism for host immune evasion by picornavirus (Dodd et al, 2001).

Mechanistically, the formation of curved Golgi cisternae leading up to its end-to-end fusion has one common denominator that cannot be ignored; all these structures have numerous NC bound to them. Further, even a solitary NC bound to Golgi membranes exhibit membrane bending. Several mechanistic hypotheses could be forwarded to explain this phenomenon, centering around the E2 glycoprotein and the NC and potentially their interaction. Work with bunyavirus glycoproteins has shown that viral glycoproteins by themselves can remodel and bend membranes and vacuolate the Golgi (Gahmberg et al, 1986). Thus, the phenomenon in alphavirus could be a result of accumulation of envelope glycoprotein and the binding of NC is just co-incidental. However, it is tempting to hypothesize that the interaction of capsid and E2 on the membrane leads to this cisternal curvature. It is also possible that the intrinsic architecture of the Golgi could be a contributor. Recent work using cryo-electron tomography has shown that the “flatness” of the Golgi cisternae is due to the presence of closely spaced yet unidentified lumenal proteins that zipper the membranes together into close apposition (18-19 nm) in the compact zone of the GA (Engel et al, 2015). The availability of the glycoproteins in the cisternal membrane leads to indiscriminate binding of the NC. Once the membrane associated E2 engages with the NC, it is possible that forces that otherwise induce budding into herniated Golgi lumen ends up bending the zippered double lamellar structure of the cisternae due to architectural constraints (Fig S9 A-F). To this end we measured and compared the lumenal width of a typical cisternae and a double lamellar CPV-II with or without detectable budding events. The results indicate that “budding like” events were always associated with increased lumenal width (herniations) indicating that the Golgi architecture may play a role in this unique phenomenology in alphaviruses (Fig S9 (C-G)). Alternatively, it could simply be that “wrapping” occurs when in situations where lipids can be replenished, whereas “budding like” phenomenon occurs when lipids are limiting in the DMV environment, leading to depletion of the inner membrane of the DMV to form SMV.

In summary, we have combined multiple innovative transmission electron microscopy (TEM) approaches to elucidate host-cell Golgi remodeling into various forms of CPV-II during VEEV infection. Using traditional thin-section TEM we first established a timeline of structural changes of the Golgi during infection where the Golgi is converted into vesicular structures. We then applied an improvised hybrid sample fixation method in conjunction with peroxide tagging of a Golgi marker to show that various forms of CPV-II in fact originate from the Golgi apparatus. To gain a three-dimensional understanding of the process we established a phased imaging approach for sequential imaging of overlapping volumes employing serial-section electron tomography. We employed the automatic image montaging capability of the Titan-KRIOS operating at 300kV at room temperature that enabled a larger volumetric coverage without compromising on the target resolution (∼4 nm). The vast array of 3D data collected enabled statistical analyses that yielded a 3D classification of the CPV-II system, identification of large and complex intermediates that links the various forms of CPV-II, relative spatial frequency of the various forms and finally a model for the morphogenesis of CPV-II. Our data identifies a second pathway of Golgi remodeling that accounts for the diverse morphological forms of CPV-II including the double lamellar form observed in alphavirus infected cells. Data from Golgi marker localization and reconstructed volume also reveal that the Golgi once converted into CPV-II forms large clusters that moves out of the perinuclear region and accumulates at the plasma membrane where it potentially transfers NC to the PM for budding. Thus, our work not only dissects a hallmark phenomenology associated with alphaviruses but also provides a path for utilizing similar resources to dissect remodeling pathways of large organelles associated in various diseases, endogenous cellular events and infections.

## Materials and Methods

### Cell culture and virus infection

Baby hamster kidney cells (BHK-21, ATCC) were cultured in Dulbecco’s modified Eagle’s medium (DMEM) supplemented with 10% fetal bovine serum (FBS) at 37°C in a 5% CO_2_ atmosphere. BHK cells were infected with the vaccine strain (TC-83) of VEEV at a multiplicity of infection (MOI) of 5, 10, 20 and 200. At various time points (3 hr, 6 hr, and 12 hr), infected cells were released from Nunc UpCell dishes non-enzymatically. Cell sheets were pelleted, resuspended in 20% BSA (Sigma-Aldrich) as a cryoprotectant, and pelleted again for cryo-fixation and preparation for electron microscopy (described below).

### Localization of horseradish peroxidase (HRP)-tagged Golgi marker, *α*-mannosidase-II

BHK cells were transfected with *α*-mannosidase-II-mCherry-HRP (Golgi FLIPPER, Kuipers et al., 2015) 8 hours prior to infection. Infection was carried out as described above, followed by pelleting and fixation with 2% glutaraldehyde for 30 min, washing 3x for 5 min with 0.1% sodium cacodylate buffer and 1x with cacodylate buffer containing 1 mg/ml 3,3’-diaminobenzidine (DAB) (Sigma-Aldrich). Pellets were then incubated for 30 min in a freshly made solution of 1 mg/ml DAB and 5.88 mM hydrogen peroxide in cacodylate buffer, pelleted and washed 3x for 5 min each in cacodylate buffer. Cell pellets were then resuspended in DMEM containing 15–20% BSA, pelleted again and cryofixed by high-pressure freezing. Samples were prepared for electron microscopy as described below. For further experimental details on the CryoAPEX method see Sengupta et al, 2019 (Sengupta et al, 2019). To determine if structural perturbation of Golgi induced by VEEV is cell type specific and to rule out artefactual effects of over expression of HRP-ManII, HeLa cells constitutively expressing the Golgi cis-medial marker α-mannosidase-II-HRP (kind gift from Dr. Franck Perez, Institut Curie) were infected with TC-83 at an MOI of 10 and 20 and cells we processed for EM identically as in the case of infected BHK cell transient expressing the HRP-ManII plasmid.

### Sample preparation for electron microscopy and tomography

Cell pellets in cryoprotectant solution were loaded onto membrane carriers (Mager Scientific, Dexter, MI) and cryo-fixed using the EM PACT2 high pressure freezer (Leica, Buffalo Grove, IL). Cryo-fixed cells were processed by freeze substitution using an AFS2 automated freeze substitution unit (Leica). Briefly, frozen pellets were incubated at -90°C in a solution of tannic acid in acetone, followed by slowly warming to room temperature in a solution of uranyl acetate and osmium tetroxide in acetone. Samples were infiltrated with Durcupan™ ACM resin (Sigma-Aldrich) and blocks were polymerized at 60°C. Thin sections (60-90 nm) were cut using the UC7 ultramicrotome (Leica), post-stained with 2% aqueous uranyl acetate and Sato’s lead, and imaged on a Tecnai T12 microscope (FEI, Hillsboro, OR) operating at 80 kV. Thicker 250-nm sections were screened on a 200 kV (CM200 Philips) microscope. Samples from all time points were screened in both thin and thick sections. For tomography, serial sections of 250-nm thickness were collected on LUXFilm coated 2 × 1 mm copper slot grids (Luxel, Friday Harbor, WA), post-stained, carbon coated, and overlaid with 10-nm colloidal gold particles (Sigma-Aldrich) for use as fiducial markers. At least 50 sections were screened for morphological determination from two representative resin blocks for each time point and condition.

### Large volume electron tomography data collection

Two large volume EM tomography data sets were collected covering an approximately 45 μm^2^ area of a BHK cell through a depth of 2 - 2.5 μm (Fig S2). Images were acquired using a Titan Krios TEM (FEI) operating at 300 kV at room temperature, outfitted with a 4k × 4k UltraScan 4000 CCD camera (Gatan, Inc., Pleasanton, CA). Tilt series were collected with automation using the program SerialEM (Mastronarde, 2003). First, a low magnification image montage of the entire grid was created to use as a map for marking regions of interest. The area of interest in the cell was visualized and marked on consecutive serial sections. At each section, a tilt series was collected. For each tilt series, images were collected at tilts from +60 to -60 degrees in 2° increments. To increase the area covered by each image, 2 × 2 montage images were collected at each tilt at 14,000x magnification, corresponding to a 0.65 nm/pixel size at the specimen level. Such montage tilt series were collected through 10 serial sections (Tomogram 1) or 7 serial sections (Tomogram 2). Smaller tomograms were collected at a pixel size of 0.8 nm, using a similar collection scheme, without serial sectioning or montaging.

### Electron tomography data reconstruction and analysis

Tilt series were aligned, and tomograms generated by weighted back projection using the eTomo interface of IMOD (Kremer et al, 1996). Serial section tomograms were aligned and joined in z to produce two reconstructed volumes with approximate dimensions 5 × 5 × 2.5 µm (Tomogram 1) and 5 × 5 × 2 µm (Tomogram 2). Membrane structures of interest were segmented by hand tracing using IMOD’s Drawing Tools and Interpolator, and 3D surface mesh models were generated. NCs were segmented by placing a sphere with a diameter of 40 nm at the center of each density. Models for the two large tomograms were joined using cellular landmarks to determine translational and rotational parameters for alignment.

Size measurements for CPV-II were obtained from the 3D mesh of the segmentation for each vesicle. The volume inside the mesh for each CPV-II was calculated in IMOD, and volume was converted to diameter assuming spherical vesicles, for ease of comparison to existing CPV-II measurements. Nearest neighbor distance analysis between each CPV-II object and each Golgi object was conducted using the *mtk* program in IMOD. Classification of NCs was achieved by analysis of the radius of curvature on adjacent membranes of Golgi cisternae and Golgi-derived vesicles. The *imodcurvature* command in IMOD was used to identify areas on the segmented model with local radius of curvature between 20 and 60 nm, corresponding to the curvature produced by a budding 40 nm NC. These areas were colored red on the membrane, as shown in Fig S6. NCs adjacent to these highly curved membranes were colored red, while NCs adjacent membranes having a radius of curvature greater than 60 nm (represented by no change in color of the membrane) were colored blue.

Details and illustration of lumenal distance measurements in Fig S9 can be found in the figure legend. At least 60 measurements were taken for each group, and the Mann-Whitney U test was used to compare each pair of distributions. Statistical analysis was done using R software (R Core Team, 2019).

## Data Availability

The manuscript does not have large-scale data sets to deposit to public databases.

## Acknowledgements

These studies were supported by the NIH (AI095366 to R.J.K. and AI081077 to R.V.S.). We thank Dr. Carolyn Machamer and Dr. Peter Hollenbeck and for their useful discussions and Dr. Saif Hassan for brainstorming sessions and help with figures. We thank Dr. Agustin Avila-Sakar and Valerie Bowman at the Purdue Cryo-EM facility for providing ready access to the Titan-KRIOS and CM200 microscopes and other instruments in the facility. We thank Dr. Christopher Gilpin at the Purdue Life Sciences Microscopy Facility for access to the T12 electron microscope. Lastly, we thank Anwesha Dasgupta for her help in editing the manuscript.

## Author Contributions

Conceptualization: R.S. and J.K.L.; Methodology: R.S., E.M.M., and J.K.L.; Formal Analysis: R.S., E.M.M., and J.K.L.; Investigation: R.S., E.M.M., S.A., and J.K.L.; Resources: J.K.L., R.J.K., and R.V.S.; Writing – Original Draft: R.S., E.M.M., and J.K.L.; Writing – Review & Editing: R.S., E.M.M., J.K.L., R.J.K., and R.V.S.; Visualization: R.S., E.M.M., and J.K.L.; Supervision: J.K.L, R.J.K., and R.V.S.; Funding Acquisition: J.K.L., R.J.K., and R.V.S.

## Supplemental figure legends

**Figure S1. Location of Arg^120^ in E2 ectodomain of VEEV-TC-83 strain**. Position 120 of the E2 ectodomain contains an arginine (Cα atom shown as red sphere; PDB ID 3J0C) in VEEV-TC-83 instead of threonine in wild type VEEV. This Thr/Arg mutation is not expected to affect the interactions of the E2 endo-domain (blue) with the capsid protein as Arg^120^ is far away from the E2 endo-domain on the opposite side of the lipid membrane (gray).

**Figure S2. VEEV infection induced progressive remodeling of the Golgi apparatus (GA) in BHK cells. (A)** Representative TEM images from 90 nm resin sections showing Golgi stacks in BHK cells infected with VEEV TC-83 at 3 hours PI. (i) Low magnification image of a cell with the perinuclear region containing Golgi stacks outlined in blue. (ii) The area from (i) with two Golgi stacks outlined in blue [magnified in (iii)] and red [magnified in (iv)]. Most stacks show mild herniations within intact stack architecture (iii) surrounded by vesicular structures [marked by red stars in (iii)]. Large herniations of cisternae were also frequently observed [yellow arrow in (iv)] alongside the rest of the normal stack [white arrow in (iv)]. BHK cells 12 hours PI exhibited disintegration of the structure of the GA (i-iv). Section from a representative cell (i), exhibited numerous Golgi stacks [at a higher magnification (ii)] that showed both extensive herniation (stack within white box) and cisternal bending and separation clearly observed at a higher magnification [magenta crescents (iii) and (iv)]. Additionally, this stage is marked by the accumulation of CPV-II vesicles containing budded virus within [(iii) and (iv), red and yellow arrowheads respectively] situated near the Golgi stacks. The size of these vesicles as well as the number of virions inside greatly varied.

The abundant accumulation of CPV-II at the perinuclear region by 6 hours PI is remarkable. To get a better idea of the spatial distribution of the CPV-II vesicles in a whole cell, 200 nm thick serial-sections were obtained, and a typical cluster of vesicles were followed through the sections **(C)**(i) sections 1 through 20). Vesicle clusters appearing from section 3 continued until section 17 indicating the large size (∼ depth of 3.6 µm) of the vesicular cluster. A representative image from the serial-section stack (section 13, (ii) and magnified images of the area demarcated within white circles (1) and (2) showed vesicular clusters enmeshed within the Golgi cisternae giving an “eggs in a nest” appearance (inset 1, blue arrow). At a higher magnification a better understanding of the morphology of the CPV-II contained within these thick sections was apparent **(D)**(i) through (x). The vesicles showed a “spiky” appearance when the surface membrane was intact indicating the presence of a dense complement of NCs on the surface [magenta arrows, (ii)]. These CPV-IIs were a mix of spherical, oblong (i, ii and iii) and even dumbbell shaped [(iv) and (v)]. A higher magnification screen of the Golgi stacks showed Golgi herniated ends of the Golgi cisternae breaking off to into dumbbell shaped elongated vesicles [(vi) area within green box magnified in (viii)]. Some cisternae were also seen to be folding upon itself into “donut-like” structures [(vi) area with blue box magnified in vii)]. Further, more complex modifications of the Golgi cisternae [(ix) and magnified image of the complex intermediate (x)] provide evidence for Golgi remodeling. Besides the perinuclear region these CPV-II clusters were also prevalent by the plasma membrane at 12 hours PI **(E)**(i) area with green box, magnified in (ii). Further magnified view of this cluster (iii) showed the presence of 4 primary classes of CPV-II [vesicles demarcated by blue boxes in (iii) and magnified view of the 4 types (1 through 4).

**Figure S3. Uninfected BHK cells exhibit canonical stacked Golgi structure**. BHK cells were processed identically as infected BHK cells (Fig S2) and exhibited well preserved cytoplasmic ultrastructure **(A-D)** including a typical stacked Golgi structure **(B-C)**. No Golgi vacuolization, fragmentation or large vesicular structures were detected around the GA as observed in infected cells.

**Figure S4. Determination of Golgi morphology in VEEV TC-83 infected HeLa cells expressing HRP-tagged-Mannosidase-II. (A)** Localization of HRP tagged Golgi resident enzyme α-mannosidase-II via TEM, shows the scattered presence of vesicular structures throughout the cytoplasm **(A)**(i) and (ii). At higher magnification, the perinuclear region [within boxed yellow region in (i)] exhibited at least two stained Golgi mini-stacks with large herniations in its cisternae surrounded by discrete stained vesicles (iii and iv). Golgi herniation is seen here to affect entire Golgi cisternae giving rise to very large vesicles (∼1-1.5 micron) (v). These stained Golgi origin vesicular structures/CPV-II are also see in small clusters at the plasma membrane (vi). It is important to note that at this late time point, the cell exhibits all stages of Golgi disintegration from herniation to vacuolization to the different morphological forms of CPV-II. **(B)** HeLa cells constitutively expressing HRP-tagged α-mannosidase-II exhibited a tight perinuclear staining like BHK cells ectopically expressing the peroxidase tagged marker. At a higher magnification, multiple stacks of a well-preserved GA were apparent **(B)**(ii) and (iii) and (v-vi). When the HeLa cell line expressing HRP tagged α-mannosidase-II was infected with VEEV/TC-83 it did not exhibit extensive scattering of the stained structures at an early timepoint **(B)**(iv), however, vacuolization of the Golgi cisternae is very apparent at 6 hours PI **(B)**(v) and (vi).

**Figure S5. Overview of large volume tomograms reconstructed using the phased collection approach. (A)** TEM image collected at 300 kV of a 250-nm thick section of a VEEV TC-83-infected BHK cell at 12 hours PI (outlined in white). The areas imaged and reconstructed are indicated by the blue and red squares. **(B-C)** A 3-nm thick virtual section from tomogram 1 **(B)** and tomogram 2 **(C)** with the z-depth along the x- and y-axis shown along the top and right side, respectively. **(D)** 3D visualization of the combined tomogram volumes with segmentation of Golgi (green), CPV-II (gold) and plasma membrane (translucent grey), and approximate tomogram boundaries in blue and red. Scale bars: **(A)** 5 μm, **(B-C)** 500 nm, **(D)** 2 μm.

**Figure S6. Color-coding of NC based on radius of curvature of the membrane to which they are bound**. The *imodcurvature* command in IMOD was used to identify areas on the segmented membrane with local radius of curvature between 20 and 60 nm, corresponding to the curvature produced by NC (radius 40 nm) in contact with the membrane. **(A)** The curled edge of a Golgi cisterna (green) with the Golgi membrane colored red in areas of radius of curvature between 20-60 nm. **(B)** The same view of the Golgi as A, without NC drawn. **(C)** NC were colored red in areas adjacent to highly curved membranes, while NC in areas with radius of curvature greater than 60 nm were colored blue.

**Figure S7. Distribution of the four classes of CPV-II**. Segmented CPV-II forms in the combined tomographic volume are shown in reference to the *trans*-most Golgi cisternae (green) and the plasma membrane (translucent grey). The distributions for class 1 **(A)**, class 2 **(B)**, class 3 **(C)** and class 4 **(D)** are shown separately. **(E)** Plot of the distance to the nearest Golgi cisternae for each CPV-II separated by class.

**Figure S8. Budding potential for NC in class 3 CPV-II assessed by available surface area**. The inner membrane surface area for class 3 CPV-II were calculated from their segmented volumes and the surface area (SA) per NC in each vesicle was determined. Percent of NC associated with membrane wrapping for each CPV-II was calculated as the fraction of inner NCs located on curved membrane, per the radius of curvature criteria as described in Fig S4. For each vesicle, the percent of NC budding was plotted against the inner membrane SA available per NC. The dashed line represents the 5000 nm^2^ per NC (conservative estimate based on a 40-nm capsid) required to wrap/envelop every NC in the vesicle. Points to the left of the line thus represent CPV-II which do not contain enough membrane surface area to fully wrap/envelop each NC they contain.

**Figure S9. The narrow lumenal width of Golgi cisternae are maintained in the bi-lamellar CPV-II and could be responsible in part for the formation of bi-lamellar CPV-II. (A-B)** Wrapping events possibly occur when multiple E2 binding domains on NC (black triangles on NC) engages the E2 glycoproteins (red) on the narrow/compact region of the GA (indicated by tightly apposed grey lines). **(C-G)** Measurement of the intralumenal distance (lumenal width) of Golgi cisternae **(C-D)** and bi-lamellar CPV-IIs **(C-E)** from 3-nm thick tomographic slices. **(C)** Tomographic slice of a region of a control unperturbed Golgi stack. **(F)** Close-up of two cisternae with red lines indicating the intralumenal distance or width of the compact region of the Golgi. **(D-F)** Intralumenal distance (red) of CPV-IIs was measured as the distance between the outer (yellow) and inner (blue) membranes (40 nm to either side of the center of an attached NC). **(G)** Comparison of intralumenal distance in intact Golgi cisterna with that of the bi-lamellar class 3 CPV-II. At least 60 measurements were taken for each of the three groups. Scale bars: 200 nm **(E)** and 100 nm **(C-F)**.

## Supplemental Video Files

**Video S1**. Overview of tomogram 1 reconstructed volume and 3D segmented structures.

**Video S2**. Golgi stack 1 and Golgi stack 2 showing slices from the tomogram transitioning into the model.

**Video S3**. Large cup-like transformation of Golgi cisterna.

**Video S4**. Overview of Golgi 1 and nearby cisternal fragments and precursors of CPVII

**Video S5**. Large pleomorphic class 1 CPV-II found in close proximity with the Golgi.

**Video S6**. Representative example of a Class 1 CPV-II.

**Video S7**. Representative example of a Class 2 CPV-II.

**Video S8**. Representative example of a Class 3 CPV-II.

**Video** S9. Representative example of a Class 4 CPV-II.

**Video** S10. 3D overview of the spatial distribution of the four classes of CPV-II in the joined tomograms

**Video** S11. Intermediate form between class 2 and 3 CPV-II containing an ostensible “fusion pore”.

**Video** S12. Intermediate form between class 3 and 4 CPV-II showing partial envelopment of several NC.

**Video** S13. Intermediate form between class 3 and 4 CPV-II showing extensive envelopment and a few fully budded virions.

**Video** S14. A CPV-II membrane in continuum with the plasma membrane depicting a state of fusion

**Video** S15. A CPV-II with a NC on its it’s outer surface bound to the inner leaflet of the plasma membrane and shows bending of this membrane exhibiting an early state of budding

