## Supplementary figures and images for "Contribution of the Golgi apparatus in morphogenesis of a virus induced cytopathic vacuolar system"

### Supplemental Figures (S1-S9)

**Figure S1**

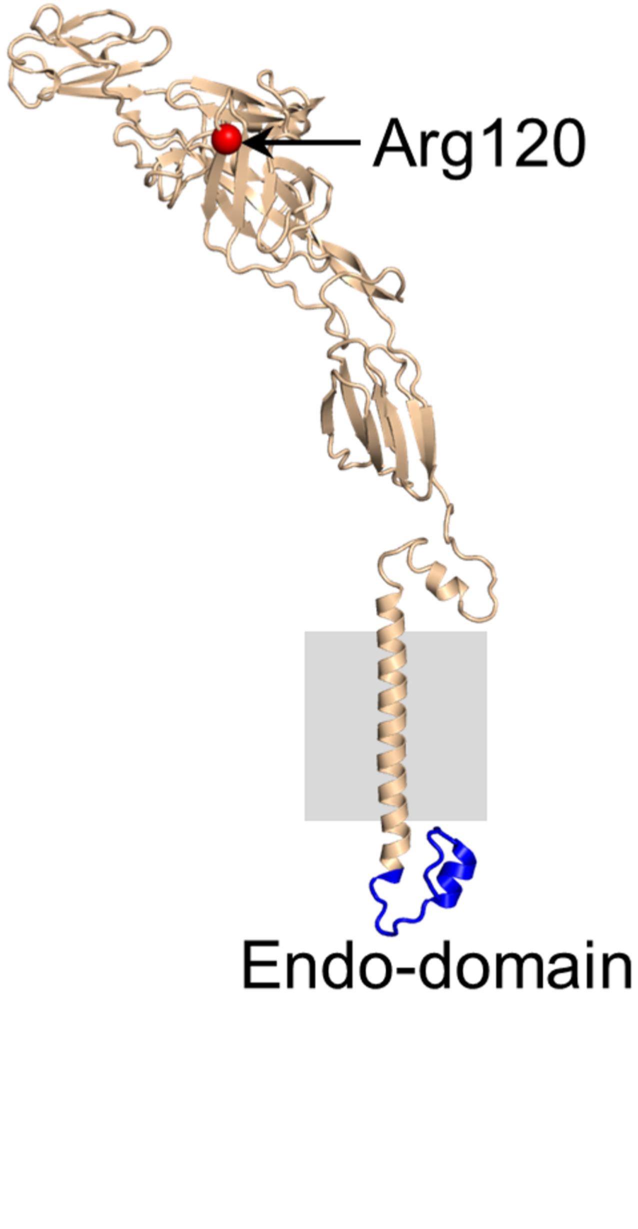

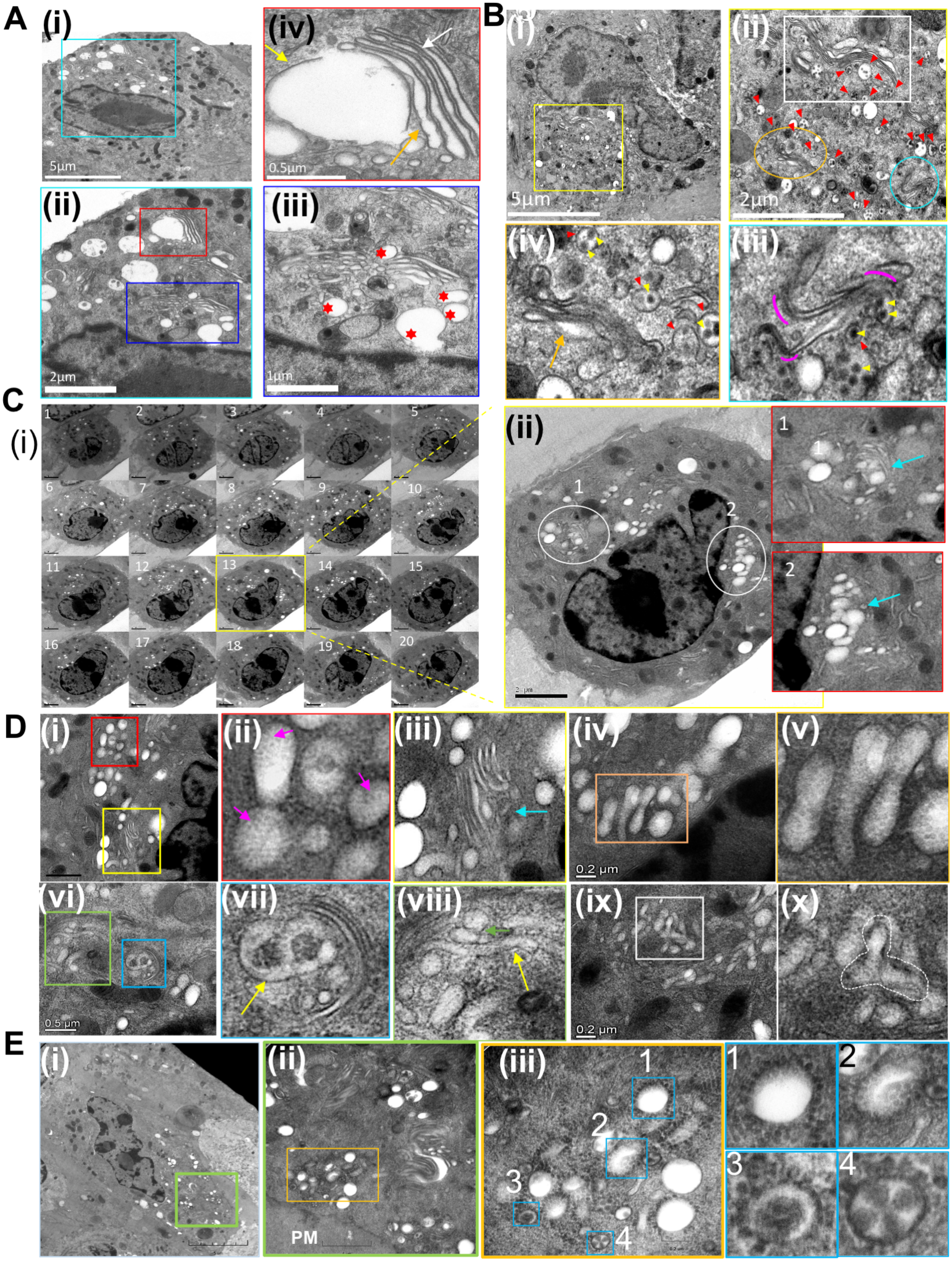

#### Figure S3

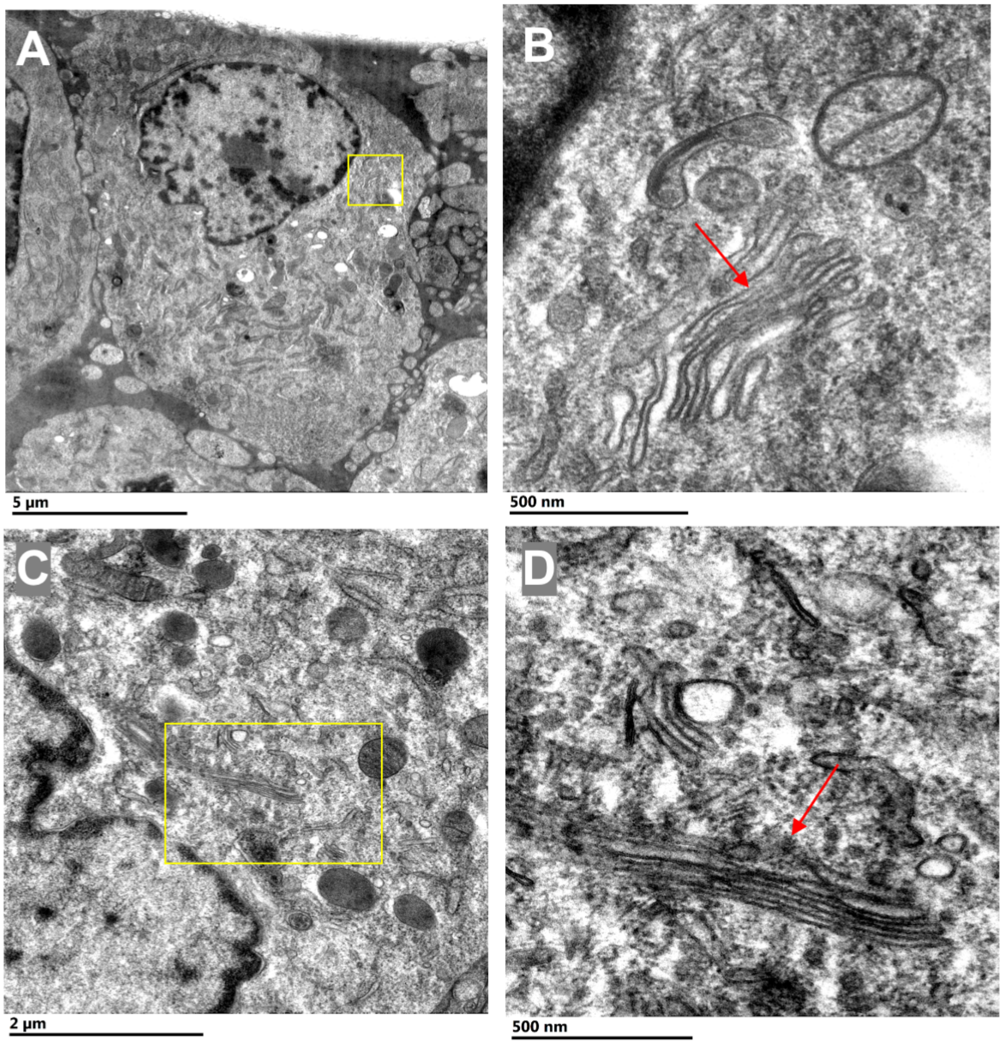

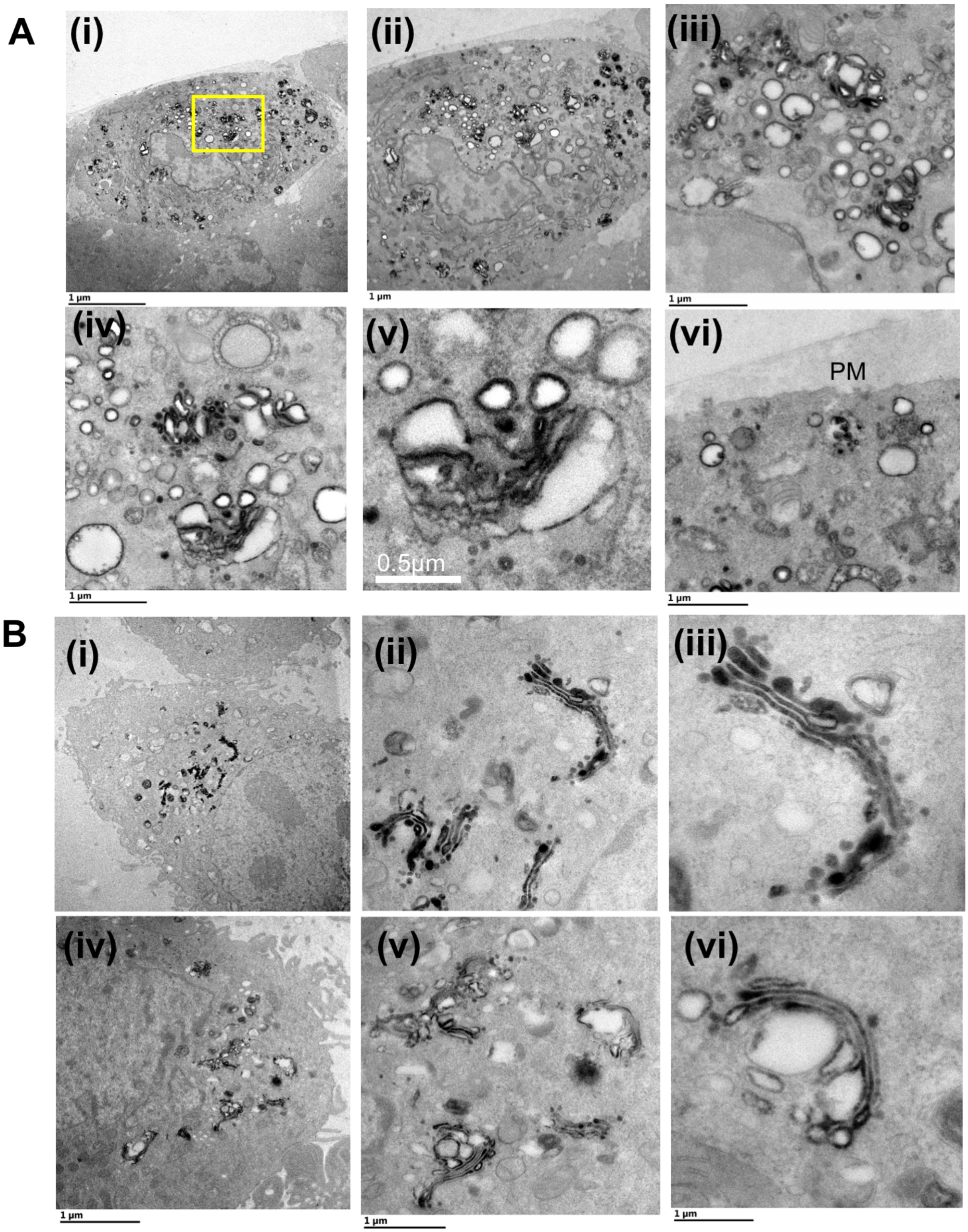

##### Figure S5

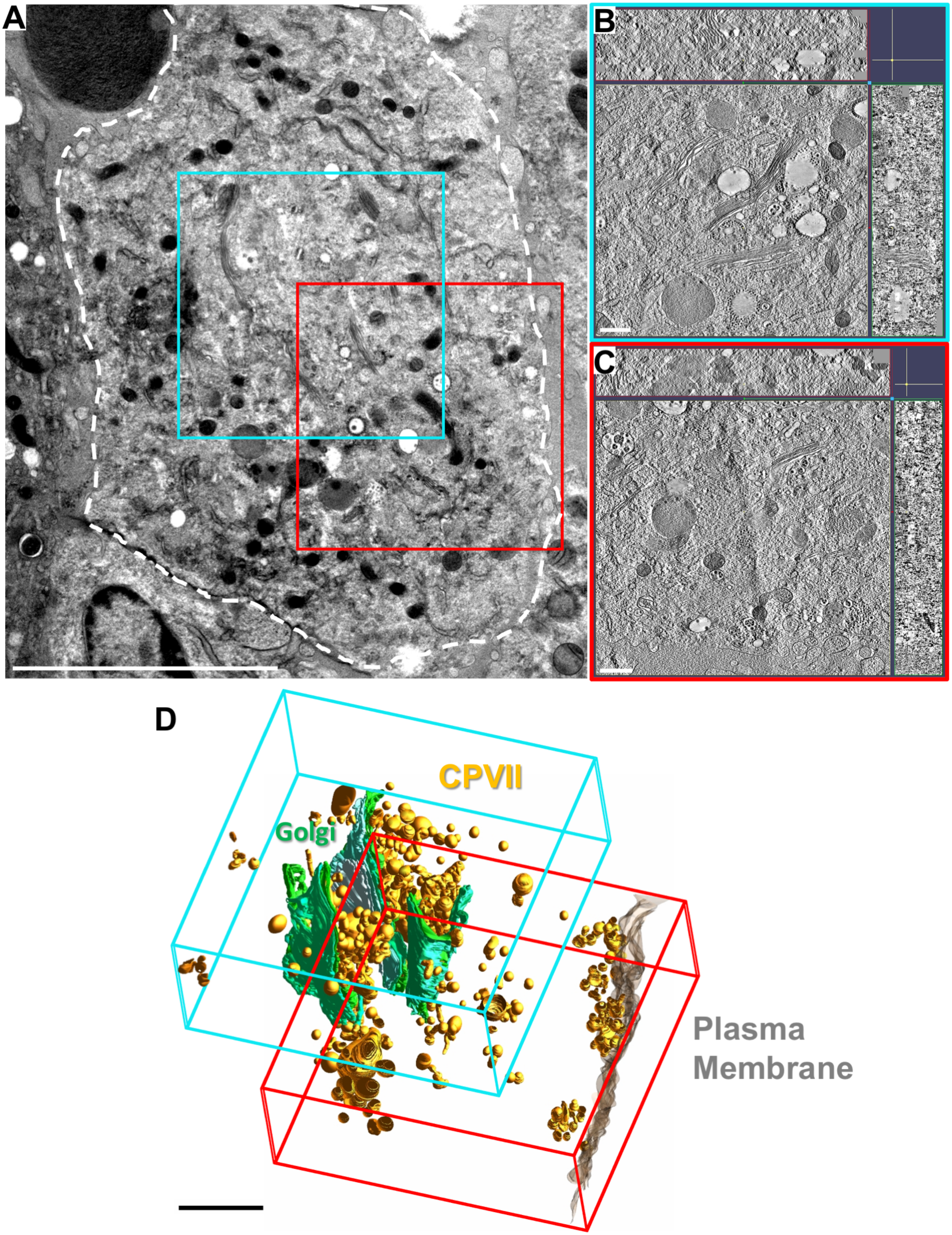

Figure S6

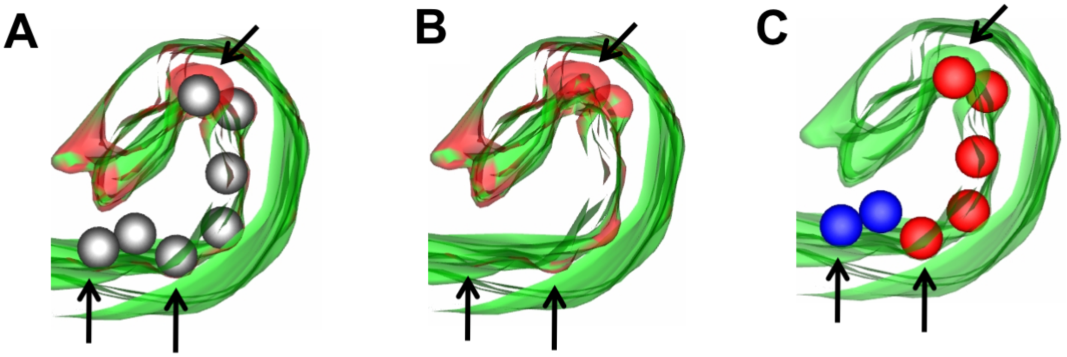

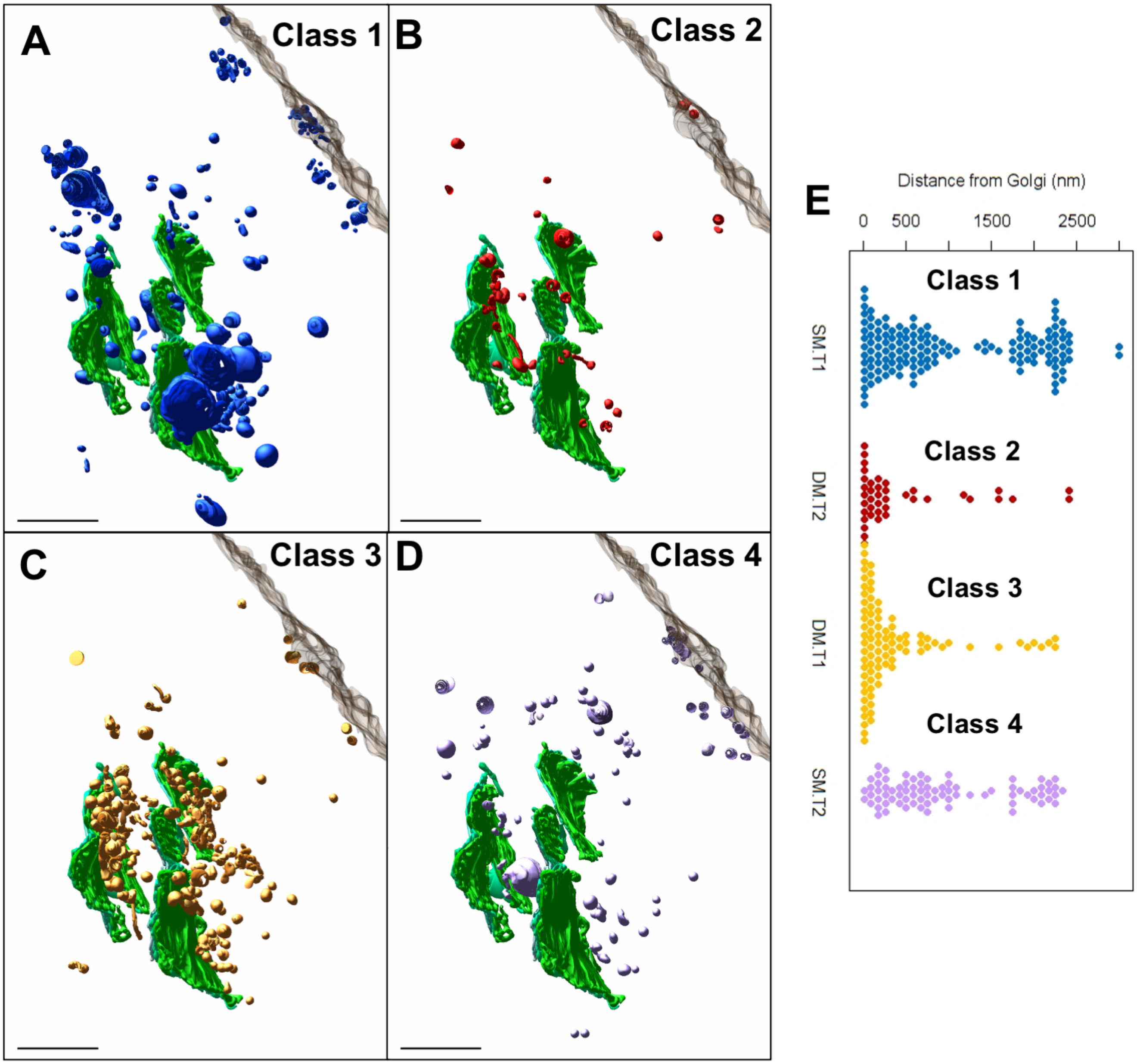

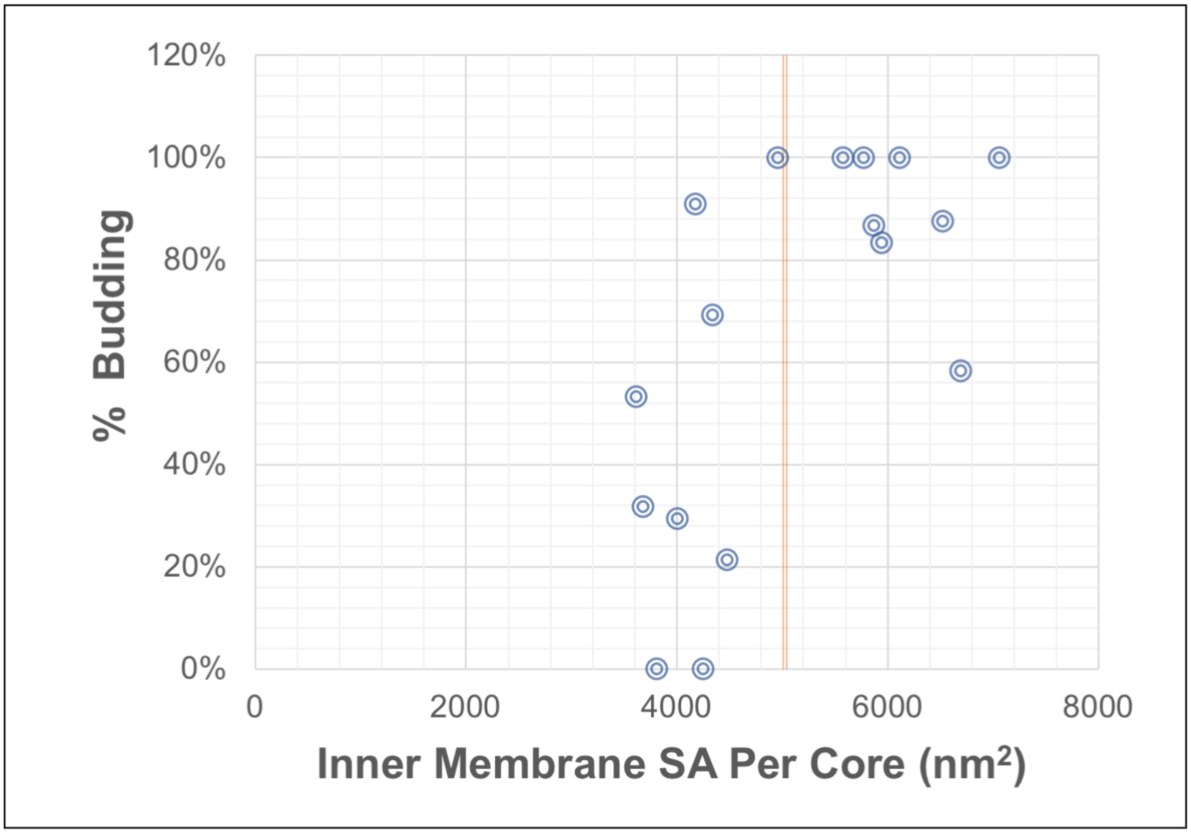

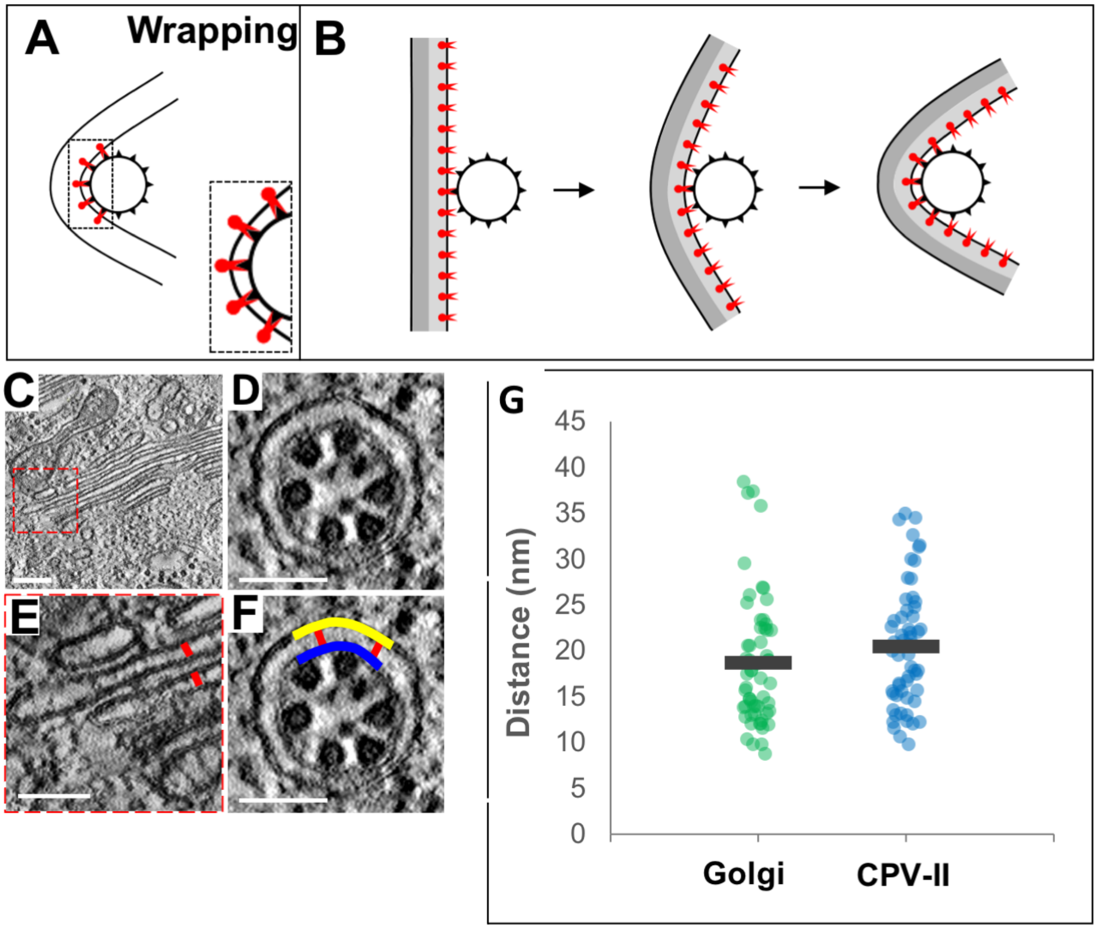
